# MreB Filaments Create Rod Shape By Aligning Along Principal Membrane Curvature

**DOI:** 10.1101/197475

**Authors:** Saman Hussain, Carl N. Wivagg, Piotr Szwedziak, Felix Wong, Kaitlin Schaefer, Thierry Izoré, Lars D. Renner, Yingjie Sun, Alexandre W. Bisson Filho, Suzanne Walker, Ariel Amir, Jan Löwe, Ethan C. Garner

**Author notes:** These authors contributed equally to this work (Saman Hussain, Carl N. Wivagg). Current affiliation: Institute of Molecular Biology and Biophysics, ETH Zürich, Switzerland. Corresponding Author: Ethan C. Garner NW 445.20, Northwest Building, 52 Oxford Street, Cambridge, MA 02138. All supplemental movies are available at http://garnerlab.fas.harvard.edu/mreb2017/.

## Abstract

MreB is essential for rod shape in many bacteria. Membrane-associated MreB filaments move around the rod circumference, helping to insert cell wall in the radial direction to reinforce rod shape. To understand how oriented MreB motion arises, we altered the shape of *Bacillus subtilis.* MreB motion is isotropic in round cells, and orientation is restored when rod shape is externally imposed. Stationary filaments orient within protoplasts, and purified MreB tubulates liposomes *in vitro,* orienting within tubes. Together, this demonstrates MreB orients along the greatest principal membrane curvature, a conclusion supported with biophysical modeling. We observed that spherical cells regenerate into rods in a local, self-reinforcing manner: rapidly propagating rods emerge from small bulges, exhibiting oriented MreB motion and increased glycan crosslinking. We propose that the coupling of MreB filament alignment to shape-reinforcing peptidoglycan synthesis creates a locally-acting, self-organizing mechanism allowing the rapid establishment and stable maintenance of emergent rod shape.

## Introduction

Although many bacteria are rod shaped, the cellular mechanisms that construct and replicate this geometry have remained largely unknown. Bacterial shape is determined by the cell wall sacculus, a giant, encapsulating macromolecule that serves to resist internal turgor pressure. One of the primary components of the cell wall is peptidoglycan (PG), which is created by the polymerization of single glycan strands linked by peptide crossbridges. Studies of isolated cell walls from rod-shaped bacteria suggest that glycan strands are generally oriented circumferentially around the rod, perpendicular to the long axis of the cell (Gan et al., 2008; Hayhurst et al., 2008; Verwer et al., 1980). This circumferential, hoop-like organization of cell wall material allows the cell wall to better resist the internal turgor pressure, as this pressure causes a stress twice as large in the circumferential direction (on the rod sidewalls) as in the axial direction (on the poles) (Amir and Nelson, 2012; Chang and Huang, 2014). This organization confers a mechanical anisotropy to the wall: the mechanically weaker crosslinks in the axial direction allows the cell wall to stretch more along its length than across its width for a given stress, and this anisotropy may assist rod-shaped cells in preferentially elongating along their length (Baskin, 2005; Chang and Huang, 2014). Concordantly, atomic force microscopy (AFM) has shown that *Escherichia coli* sacculi are 2-3 times more elastic along their length than across their width (Yao et al., 1999). This rod-reinforcing circumferential organization is also observed in the cell walls of plants; hypocotyl and root axis cells rapidly elongate as rods by depositing cellulose fibrils in circumferential bands around their width, resulting not only in a similar dispersive rod-like growth, but also a similar anisotropic response to stress (Baskin, 2005). The organized deposition of cellulose arises from cortical microtubules self-organizing into a radial array oriented around the rod width, and this orients the directional motions of the cellulose synthases to insert material in circumferential bands (Bringmann et al., 2012; Paredez et al., 2006).

In contrast to our understanding of the self-organization underlying rod-shaped growth in plants, how bacteria construct a circumferential organization of glycan strands is not known. This organization may arise via the actions of a small number of genes known to be essential for the formation and maintenance of rod shape. Collectively termed the Rod complex, these include the conserved *mreBCD* operon (Wachi and Matsuhashi, 1989) and the glycosyltransferase/transpeptidase enzyme pair RodA/Pbp2 (Cho et al., 2016). The spatial coordination of RodA/Pbp2-mediated PG synthesis is conferred by *mreB*, an actin homolog (Jones et al., 2001; van den Ent et al., 2001). MreB polymerizes onto membranes as antiparallel double filaments, which have been observed to bend liposome membranes inward (Fig. 1A) (Salje et al., 2011; van den Ent et al., 2014). Loss or depolymerization of MreB causes rod-shaped cells to grow as spheres (Jones et al., 2001). *In vivo*, MreB filaments move circumferentially around the width of the rod (Domínguez-Escobar et al., 2011; Garner et al., 2011; van Teeffelen et al., 2011). Super-resolution imaging has demonstrated that MreB filaments always translocate along their length, moving in the direction of their orientation (Olshausen et al., 2013). MreB filaments move in concert with MreC, MreD, and RodA/Pbp2 (Domínguez-Escobar et al., 2011; Garner et al., 2011), and loss of any one component stops the motion of the others. The directional motion of MreB filaments and associated Rod complexes depends on, and thus likely reflects, the insertion of new cell wall, as this motion halts upon the addition of cell wall synthesis-inhibiting antibiotics (Domínguez-Escobar et al., 2011; Garner et al., 2011; van Teeffelen et al., 2011), or specific inactivation or depletion of Pbp2 (Garner et al., 2011; van Teeffelen et al., 2011) or RodA (Cho et al., 2016).

**Figure 1.**
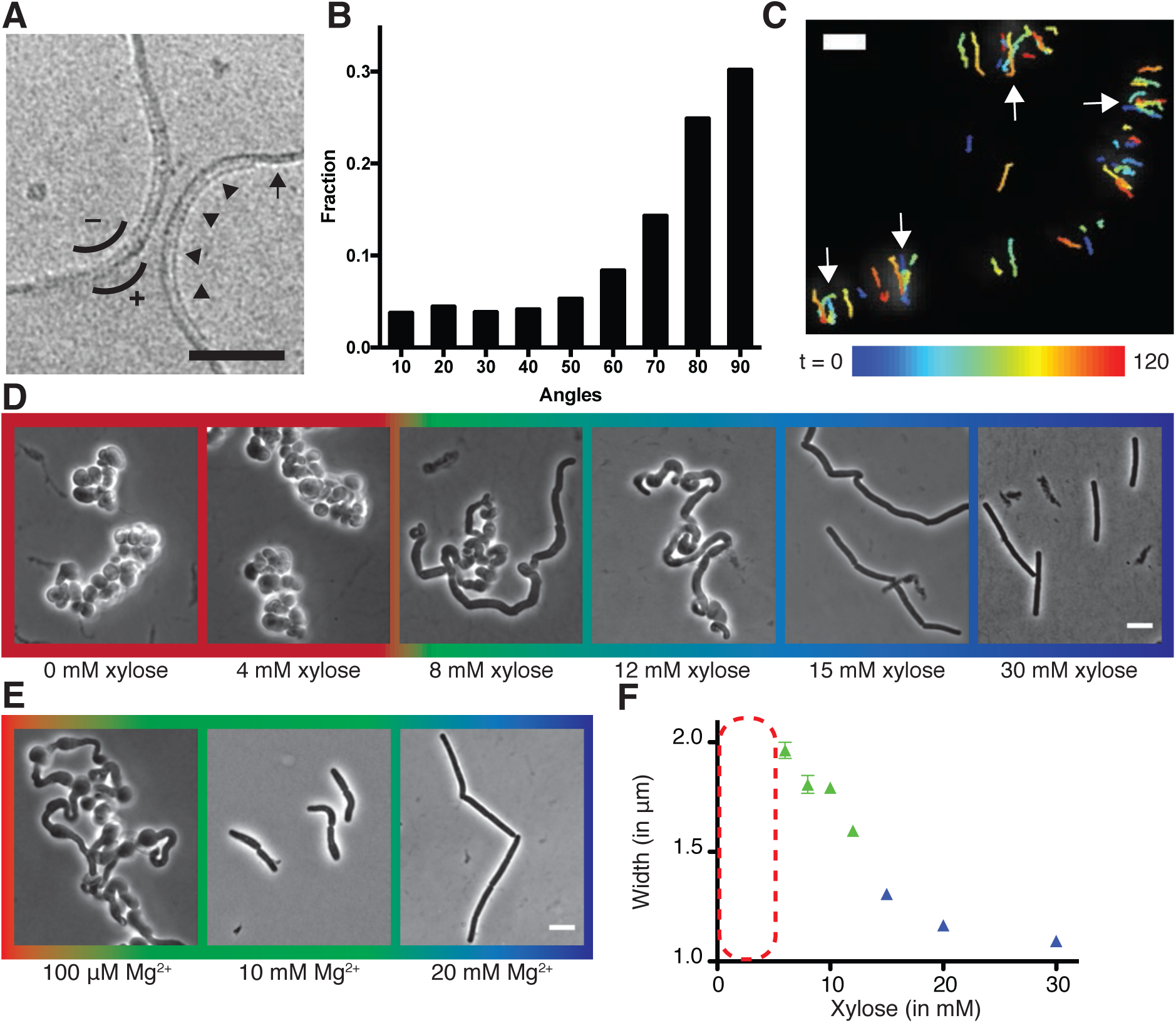
Curved MreB filament motions do not follow an ordered template (A-C). **(A)** The negative curvature of MreB filaments (arrowheads) aligns with the negative principal curvature of the liposome surface (arrow). Scale bar is 50 nm. **(B)** Angular distribution of GFP-Mbl trajectories relative to the long axis of the cell indicates that while the distribution has a mode of 90°, it is broad (SD = 34°). **(C)** Particle tracking of Mbl-GFP during 100 seconds (1 rotation) indicates trajectories close in time frequently cross paths (white arrows). Scale bar is 1 μm. **Modulating teichoic acid levels titrates cell width and shape (D-F). (D)** Strains with *tagO* under inducible control display a teichoic acid-dependent decrease in width. **(E)** BEG300 at an intermediate level of *tagO* induction (15mM xylose) shows a Mg^2+^ dependent decrease in width. All scale bars are 5 μm. See also Figure S1. **(F)** Plot of cell width as a function of *tagO* induction in LB supplemented with 20 mM Mg^2+^, calculated from rod-shaped cells (error bars are Standard Error of the Mean (SEM)). Areas not plotted at lower xylose levels (red dashed rectangle) are regions where cells are round (no width axis). Color scheme for D-F: red indicates round cells (no width axis), blue indicates rods (measurable width axis), and green indicates intermediate regimes where both rods and round cells are observed.

It is not known how MreB and its associated PG-synthetic enzymes construct rod-shaped cells. As the motions of the Rod complexes reflect the insertion of new cell wall, their circumferential motions could deposit glycans in the hoop-like organization required to both build and reinforce rod shape. Therefore, we worked to understand the origin of this circumferential organization, seeking to determine what orientsμ the motions of MreB and associated enzymes around the rod width in *Bacillus subtilis*. *B. subtilis* contains 3 MreB paralogs (MreB, Mbl, and MreBH) that co-polymerize into mixed filaments and always colocalize *in vivo* (Defeu Soufo and Graumann, 2006) (Soufo and Graumann, 2010) (Dempwolff et al., 2011). Thus, we assume throughout that MreB and Mbl are interchangeable for fluorescent imaging.

## Results

### Oriented MreB Motion Cannot Arise from an Ordered Cell Wall Template

The mechanism by which MreB filaments and associated PG synthases orient their motion around the rod circumference is not known. Each filament-synthase complex is disconnected from the others, moving independently of proximal neighbors (Garner et al., 2011). The organized, circumferential motion of these independent filament-synthase complexes could arise in 2 ways: 1) A templated organization, where cell wall synthetic complexes move along an existing pattern of ordered glycan strands in the cell wall as they insert new material into it (Holtje, 1998), or 2) A template-independent organization, where each synthetic complex has an intrinsic mechanism that orients its motion and resultant PG synthesis around the rod circumference.

To explore the extent of order within the motions of the Rod complex, we analyzed the trajectories of GFP-Mbl filaments during the period of one axial revolution using total internal reflection fluorescence microscopy (TIRFM). We observed that filament trajectories close in time (within the period of one revolution) frequently cross (Fig. 1C, Movie S1), making it unlikely that MreB filaments move along a perfectly ordered template. Rather, the motions of filaments are overall oriented, but not perfectly aligned, a characteristic reflected by the broad distribution of angles that MreB (Fig. 1B) and the other components of the Rod complex move relative to the long axis of the cell (Domínguez-Escobar et al., 2011; Garner et al., 2011). As MreB movement reflects the insertion of new glycan strands, these motions indicate that the sacculus is built from somewhat disorganized, yet predominantly circumferential strands, a conclusion in agreement with previous studies that assayed cell wall organization with cryo-electron microscopy (Gan et al., 2008), atomic force microscopy (Hayhurst et al., 2008), and X-ray diffraction (Balyuzi et al., 1972). Furthermore, preexisting cell wall is not necessary for the regeneration of rod shape from wall-less *B. subtilis* L-forms (Kawai et al., 2014), indicating that both oriented MreB motion and rod shape can arise without an ordered template.

### MreB Motions Become Isotropic in the Absence of Rod Shape

As it appeared that organized MreB motion does not arise from patterns in the cell wall, we hypothesized there was an intrinsic mechanism orienting the motion of each MreB filament-cell wall synthetic complex. To test this hypothesis, we examined MreB motions as we changed the shape of cells from rods to spheres. As the internal turgor pressure and stiffness of *B. subtilis* resists external mechanical perturbations to its shape (Renner et al., 2013), we first altered the shape of cells by controlling the level of wall teichoic acids (WTAs). WTAs are negatively charged cell wall polymers believed to increase the rigidity of the sacculus via their coordination of extracellular Mg^2+^ (Matias and Beveridge, 2005) or modulation of hydrolase activity (Atilano et al., 2010). Knockouts in *tagO*, the first gene in the WTA synthesis pathway, create large, slow-growing, round cells that still synthesize PG, building extremely thick and irregular cell walls (D’Elia et al., 2006). We placed *tagO* under xylose-inducible control and grew cells at different induction levels. At high TagO inductions, cells displayed normal widths, as expected. As we reduced TagO levels, rods became gradually wider (Fig. 1D,1F) until, beneath a given induction, cells were no longer able to maintain rod shape, growing as spheres (or clumps of spheres) with no identifiable long axis. At intermediate induction levels, we observed a transition region between the two states, with cells growing as steady state populations of interconnected rods and spheres (Fig 1D). In agreement with models that cell wall rigidity is conferred via WTA-mediated coordination of Mg^2+^ (Thomas and Rice, 2014), both cell width and the amount of TagO induction determining the rod/sphere transition could be modulated by Mg^2+^ levels (Fig. 1E, S1).

By tracking the motion of GFP-MreB filaments in these differing cell shapes, we found that motion is always oriented in rods, moving predominantly circumferentially at all induction levels above the rod/sphere transition. However, in round cells (those induced beneath the rod/sphere transition point or in *tagO* knockouts) MreB filaments continued to move directionally, but their motions were isotropic, moving in all directions (Fig. 2A, Movie S2A). To quantify the relative alignment of MreB under each condition, we calculated the angle between trajectory pairs less than 1μm apart (Fig. 2B, S2A). This analysis revealed that MreB motions are more aligned when cells are rods: above the rod/sphere transition, trajectories have a median angle difference of 26°; while at low TagO inductions, where cells are round, the angle difference increases to 42°, close to that of randomly oriented trajectories (45°).

**Figure 2.**
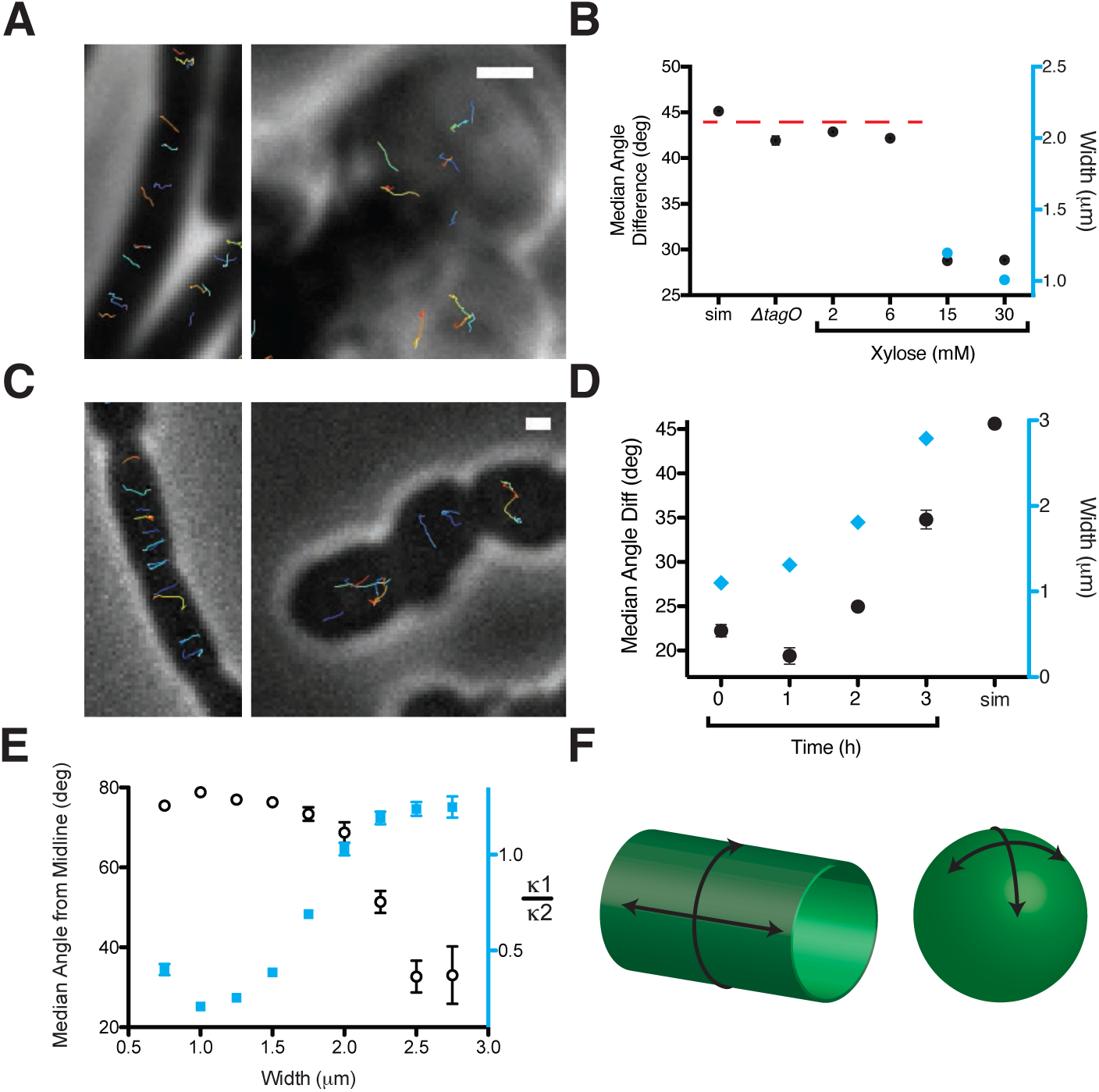
Oriented MreB motion correlates with rod shape. **(A)** BEG300 at maximum *tagO* induction (30 mM) is rod-shaped, and MreB tracks are largely oriented perpendicular to the midline of the cell *(left)*. Δ*tagO* cells show round morphologies with unaligned MreB motion (*right*). **(B)** Median inter-track angle difference for track pairs ≤ 1 μm apart, plotted for BEG300 at several *tagO* induction levels, Δ*tagO* cells, and a simulation of randomly oriented angles (*sim*).

To verify that the loss of oriented MreB motion was due to the changes in cell shape, and not from some other effect of reduced WTA levels, we created round cells by alternate means. Depletion of both elongation PG transpeptidases (Pbp2a and PbpH) causes rod-shaped cells to become wider over time as they convert to spheres (Garner et al., 2011). We used this gradual transition of rods into spheres to examine both the width and overall shape dependence of MreB motion. At initial points of depletion (1 - 2 hours) the rods widened but maintained circumferential MreB motion. At 2.5 hours of PbpA depletion, cells were a mix of spheres and rods of differing widths. These cells displayed the same pattern of MreB orientation observed above: round cells contained unoriented MreB, while nearby rod shaped cells showed circumferential motion (Fig. 2C, Movie S2B). Quantitation of trajectories from all cells (both rods and spheres) at each time point of depletion indicated an increase in the median angle between trajectories as the population grew wider and rounder over time (Fig. 2D, S2D).

In *E. coli,* the angle of mutant MreB filaments relative to the long axis has been reported to increase with cell width (Ouzounov et al., 2016). To test if the angle of MreB movement changes with respect to cell width in *B. subtilis*, we calculated the angle of each trajectory to the long axis for all cells in our data with an identifiable width axis. At the same time, we also measured the curvature of each cell to determine how the overall shape of the cell affected the orientation of motion (Fig. S2E). This revealed that MreB motion in rods remained equivalently oriented over a wide range of rod widths, up to ∼2 μm (Fig. 2E, S2B, S2C). Beyond a 2 μm width, cells began to lose their rod shape as they became more spherical, and the predominantly circumferential orientation of MreB motion was lost (Fig. 2E, S2E). This suggested that oriented MreB motion does not sense or rely on a specific cell radius; rather the orientation relies on differences between the two principal curvatures of the membrane. It appears that the motion of MreB filaments is oriented along the direction of greatest principal curvature: In rods, there is zero curvature along the rod length, and high curvature around the rod circumference, along which filaments orient. In contrast, in round cells where MreB motion is isotropic, the two principal curvatures are equal (Fig. 2F).

For spherical cells width is not measurable, indicated with a dashed red line. **(C)** *ΔpbpH* cells with *pbpA* under IPTG control display aligned MreB motion when *pbpA* is fully induced and cells are rods (*left*), but display unaligned MreB motion as Pbp2a levels reduce and cells become round (*right*). **(D)** Median inter-track angle difference for track pairs 1 μm apart during Pbp2a depletion with cell widths at each time point. **(E)** Median angle from the midline (white circles) calculated for all rod-shaped cells from experiments in 2A-D plotted as a function of cell width. MreB filament alignment falls off rapidly beyond 2 μm, a point corresponding to where cells become round, as shown by the ratio of principal curvatures (blue squares) approaching 1. See Fig S2E for further explanation. **(F)** Schematic showing the difference between the 2D surface curvature profile of rods and spheres. On the inside surface of spheres, all points have negative, yet equal values for both principal curvatures. In rods, however, one principal curvature is negative (the radius), while the other is 0 (the flat axis along the rod). All scale bars are 1μm. All error bars are SEM. See also Figure S2.

### MreB Aligns Within Round Cells and Protoplasts Forced into Rod Shape

To further verify that MreB filaments orient in response to overall cell shape, we externally imposed rod shape on cells with unoriented MreB motion. We loaded TagO*-*induced cells into long 1.5 ×1.5 μm microfluidic chambers, then reduced TagO expression to levels insufficient to produce rods in liquid culture (Fig. 3A, S3A). After TagO depletion, cells expanded to fill the chamber indicating that WTA-depletion caused shape changes just as in bulk culture (Fig. 3A, S3A). Within these chambers, cells grew as rods, but at a wider width (1.5 μm) than wild-type cells, set by the chamber. When cells grew out of the chamber they swelled just as in bulk culture, showing confinement was required for rod shape at this induction level (Fig. 3B, S3A). In the TagO-depleted cells confined into rod shapes, MreB moved circumferentially (Fig. 3C, Movie S3) with an angular distribution similar to that of wild-type cells (90°, SD = 36 °) (Fig. 3F(i)), confirming that MreB orients in response to the cells having rod shape. This experiment demonstrates that the isotropic MreB motion observed in round cells arises from the lack of rod shape, and not from some other effect of our genetic perturbations. This experiment also showed another unexpected result: the doubling time of free (unconstrained) cells induced at similar TagO levels is slow (53 ± 10 min), but confining them into rod shape restored their doubling time (44 ± 4 min) toward wild-type rates (39 ± 9 min) (Figure S3C).

**Figure 3.**
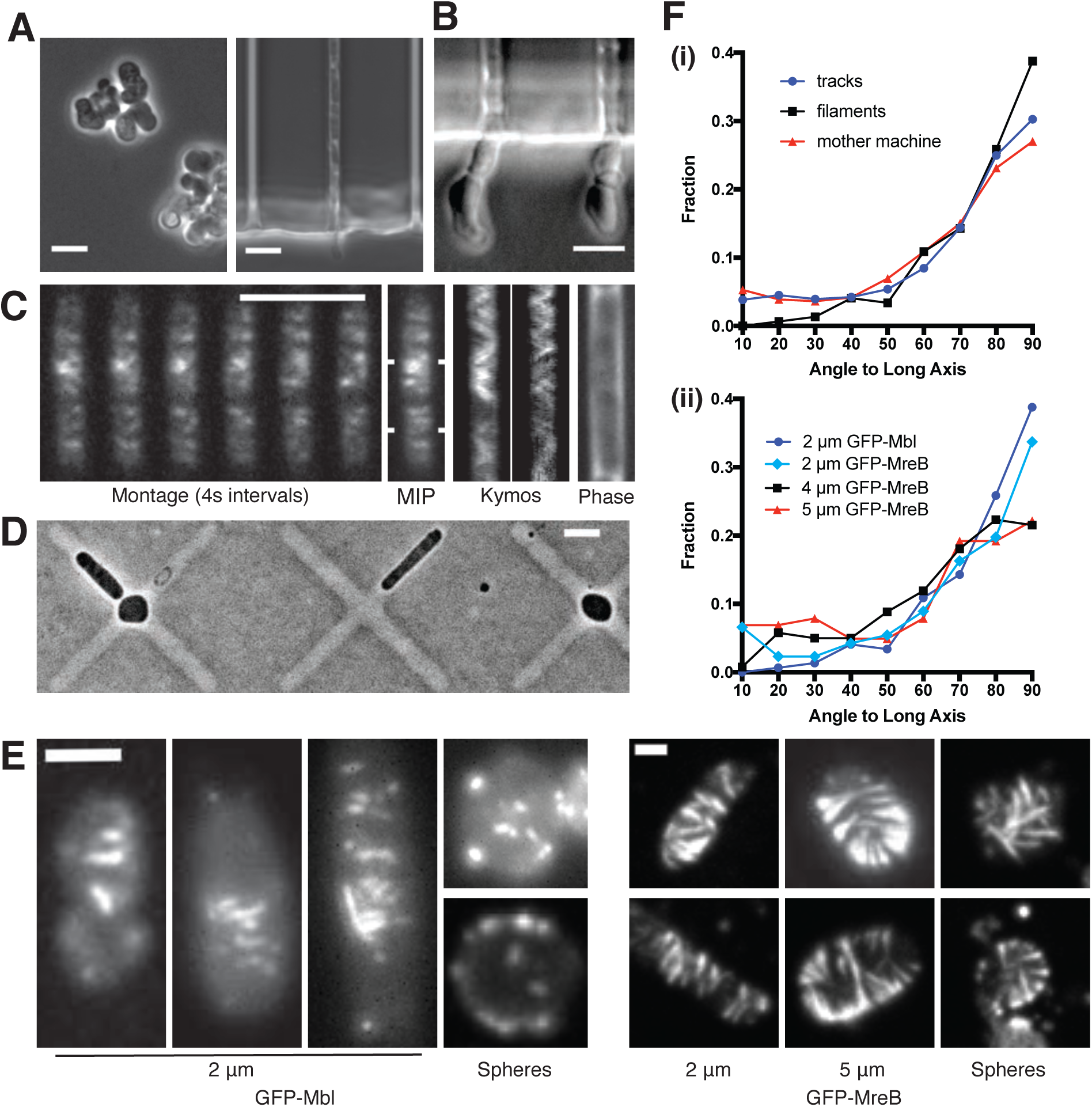
MreB filaments orient when rod shape is induced by external confinement. **(A)** Phase contrast images of BEG300 grown in LB supplemented with 2 mM xylose and 20 mM Mg^2+^ in bulk culture (*left*) or confined into microfluidic channels of 1.5 ×1.5 μm (*right*). **(B)** Confined cells induced at 3 mM xylose in 20 mM Mg^2+^ progressively swell upon escaping confinement into free culture. See also Figure S3A. **(C)** (*Left*) Montage of MreB filaments moving across a confined cell. (*Right*) Maximal intensity projection of montage, kymographs of marked points and a phase contrast image of the cell. Scale bars for a-c = 5 μm. **(D)** Phase contrast images of protoplasts contained in agar crosses. Cells in the center grow to be round while cells in arms grow as elongated rods. **(E)** (*left)* Short GFP-Mbl filaments orient circumferentially in rod-shaped protoplasts (*2 μm*) but lack orientation in round protoplasts *(spheres)*. *(right)* Long GFP-MreB filaments orient in rod-shaped protoplasts (*2 μm*); GFP-MreB filaments are still oriented in wider rod-shaped protoplasts (*5 μm*), but not to the same extent. In round protoplasts, GFP-MreB filaments are unoriented *(spheres).* See also Fig. S3B. Scale bar is 2 μm. **(F) (i)**The angular distribution of filaments within protoplasts is centered at 90° (SD 25°, n=147), similar to that of MreB motion in TagO-depleted, confined cells (90°, SD 36°, n=359) and MreB motion in wild-type cells (88°, SD 34°, n=1041). **(ii)** In channels of varying widths (2, 4 and 5 μm), the orientation of GFP-MreB filaments remains circumferential but the angular distribution becomes wider at increasing channel width (93°, SD 34°, n=258 at 2 μm), (81°, SD 35°, n=260 at 4 μm) and (86°, SD 41°, n=203 at 5 μm).

Next, we attempted to decouple MreB filament orientation from both A) the directional motion of filaments, and B) any structure within the cell wall. To accomplish this, we examined filament orientation in protoplasts (cells without cell wall) that we confined into different shapes, using highly expressed GFP-MreB to assay long filaments, and GFP-Mbl to assay short filaments. We protoplasted cells in osmotically stabilized media (Wyrick and Rogers, 1973), then grew them under agar pads containing micro-patterned cross shapes. Cells in the center of these crosses (∼5 μm diameter) were forced to grow as spheres, whereas cells in the arms were constrained to grow into rods of various widths ranging from 2-5 μm. (Fig. 3D). Cells growing in these molds did not produce cell wall, as determined by WGA staining (Fig. S3B). As reported previously (Domínguez-Escobar et al., 2011), MreB filaments within protoplasts did not move directionally (Movie S4), likely because the cell wall provides the fixed surface across which the PG synthesis enzymes move. Within the protoplasts confined into the smallest rod shapes (2 μm), filaments oriented at a distribution of angles predominantly perpendicular to the cell length (Fig. 3E-F). The angular distributions of short GFP-Mbl filaments and longer GFP-MreB filaments were similar to each other (94**°** ± 25 and 93° ± 34 respectively_),_ and also similar to the distribution of filament trajectories observed in intact, wild-type cells (88**°**, SD = 34**°**). As we increased the width of the imposed rod shape from 2 to 5 μm, filaments remained predominantly oriented in all cases, but the distribution of alignment became increasingly broad (86**°**, SD = 41**°** at 5 μm). In contrast to confinement in rods, both short and long filaments in spherically confined protoplasts remained unoriented (Fig. 3E). Together, these data demonstrate that MreB filaments orient to point around the rod width even in the absence of cell wall or directional motion, as long as the cell has a rod shape. These experiments also demonstrate that MreB filaments will align even in wider rods, where the difference in principal curvatures is smaller than in wild-type cells, but that, as the difference in principal curvatures decreases, filament alignment becomes more disordered.

### MreB Filaments Orient Around Liposome Tubes *in vitro*

To test if MreB filaments are themselves sufficient to align along the predominant direction of membrane curvature, we assembled purified *T. maritima* MreB within liposomes and visualized it using electron cryo-electron microscopy and tomography. While controlling the final concentration of protein encapsulated within liposomes ≤ 1μm is difficult, we were able to assemble MreB inside liposomes at high concentrations. At these concentrations, MreB filaments tubulated liposomes, creating rod-like shapes (Fig. 4A, S4, Movie S5). In tubulated regions, MreB filaments could be traced around the circumference of the liposome tube, while filaments in spherical regions were found in all possible orientations (Fig. 4A). At the highest concentrations, tubulated liposomes contained closely packed filament bundles, allowing us to observe a regular patterning of the canonical double filaments of MreB (Fig. 4B). Purified wild-type MreB did not bind to the outside surface of small liposomes contained within larger ones (Fig. 4A), indicating that MreB filaments preferentially polymerize on inward (negative) curvatures, akin to the inner leaflet of the bacterial membrane. In the absence of MreB, liposomes are spherical, with no deformations (Fig. S4C). Together, this data suggests that MreB filaments themselves are sufficient to align along the predominant direction of membrane curvature, as observed here with laterally associated filaments. We note that the experimental limitations of the liposomal system, combined with the tendency of MreB filaments to self-associate make it difficult for us to acquire and study the alignment of single filaments *in vitro*. Also, it remains to be determined if membrane-associated MreB filaments exist as bundles or isolated filaments *in vivo.*

**Figure 4.**
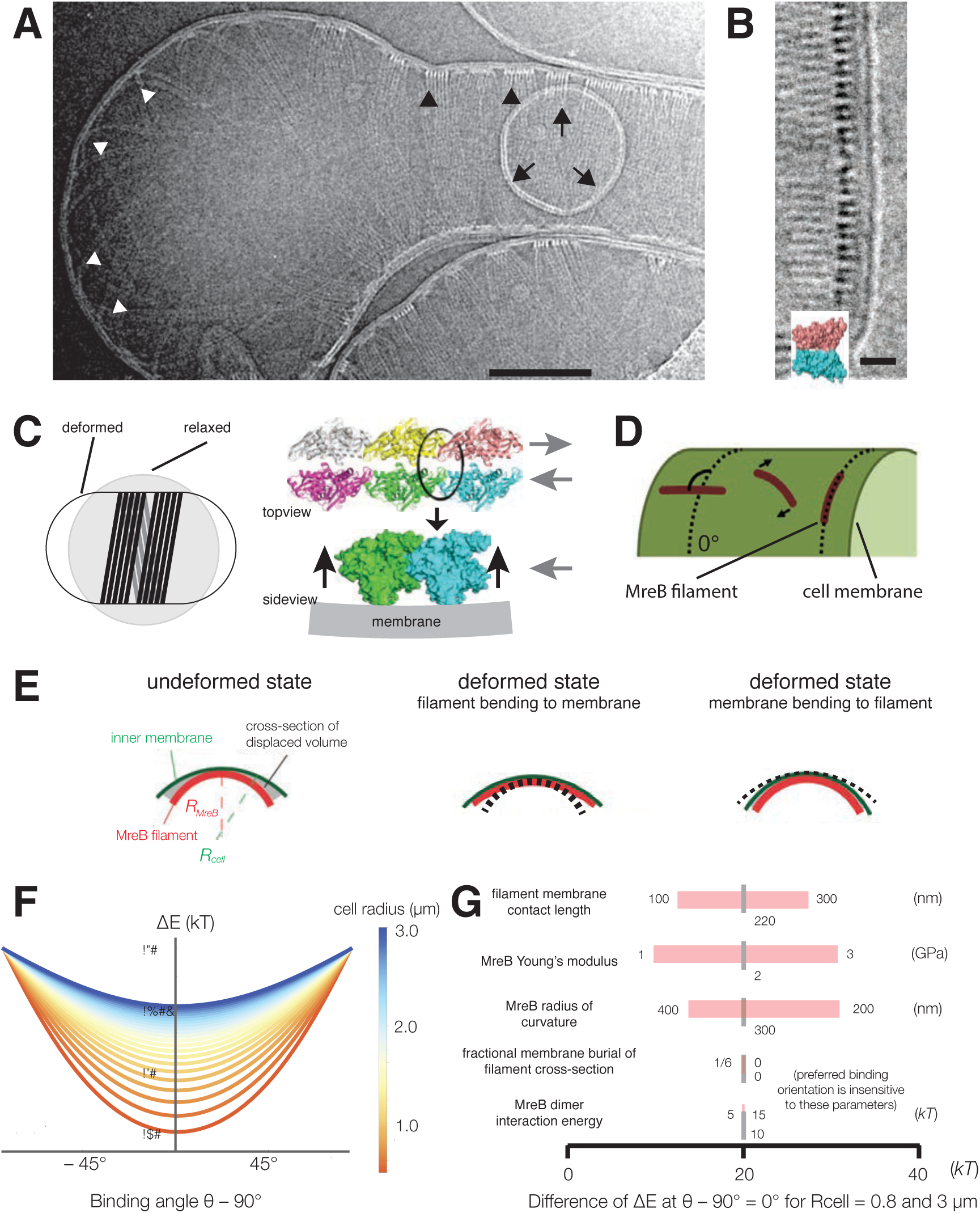
*T. maritima* MreB filaments assembled in liposomes align perpendicular to the rod axis. **(A)** Black arrowheads show aligned bundles of filaments in a tubulated liposome, white arrowheads show unaligned bundles in a spherical region of the same liposome. Arrows show a positively curved surface inside the liposome, to which no MreB filaments bind. Scale bar is 100 nm. See also Figure S4. **(B)** MreB in liposomes adopts a double-stranded antiparallel protofilament arrangement consistent with (van den Ent et al., 2014). Scale bar is 50 nm. **(C)** (*left*) Schematic drawing depicting the cause of the shape change from spherical to rod-shaped liposomes: MreB wants to attain greater curvature and since there are many filaments, they are laterally stabilized. As the liposome is much more easily deformable than cells, the resulting energy minimum is a deformed liposome with an MreB helix on the inside. (*right*) Model showing why the unusual architecture of MreB filaments might have been selected during evolution: its juxtaposed subunits in the two antiparallel protofilaments produce putative hinges that could be the region of bending for these filaments. Canonical F-actin filament architectures, with staggered subunits, would need bending within the subunits, which is less easily achieved. **Modeling of MreB – membrane interactions and filament orientation. (D, E)** Hydrophobic residues are located on the outer edge of the antiparallel MreB double filament, which is here modeled as an elastic cylindrical rod. To achieve maximum hydrophobic burial, membrane deformation, MreB bending, or a combination of the two may occur. **(F)** A plot of the change in total energy (ΔE) caused by the MreB-membrane interaction against the binding angle θ for various cell radii shown in the color scheme on the right. Note that ΔE is minimal at θ=90°, which agrees with the observed orientation of MreB binding and motion. At larger rod radii, the energetic well becomes flatter and MreB binding becomes more susceptible to thermal fluctuations and other sources of stochasticity, which would result in a broader angular distribution of filaments. **(G)** A sensitivity analysis of the model over a range of model parameters.

### Biophysical Modeling Suggests Highly Bent MreB Filaments Orient Along the Greatest Principal Curvature to Maximize Membrane Interactions, a Prediction Insensitive to Large Variations in Parameters

The above observations demonstrate that MreB filaments sense and align along the direction of greatest principal curvature, i.e., the more curved inner surface of the rod circumference. The ultrastructure of MreB filaments provides a possible mechanism: MreB filaments are bent (Salje et al., 2011), with the membrane-interacting surface on the outer face of the bend (Fig. 4C). This bent conformation could cause filaments to preferentially orient along the curved rod circumference, rather than the flat rod length, to maximize the burial of hydrophobic moieties into the membrane, a mechanism suggested by previous theory (Wang and Wingreen, 2013).

As the curvature of MreB filaments bound to liposomes is much greater (∼200 nm diameter) than that of *B. subtilis* cells (∼900 nm diameter), we performed analytical calculations to model how highly curved MreB filaments would align within a cell with a less curved surface (Fig. 4D-G, Supplemental Text 1). As many of the biochemical and physical parameters of MreB are still unknown, we first assumed a fixed set of parameters, and later verified that our results were robust over a large parameter range. We initially assumed a membrane interaction energy of 10 kT per monomer (calculated from residues involved in membrane associations (Salje et al., 2011)), and a similar Young’s modulus to actin (2 GPa). We modeled filaments as elastic beams made of two protofilaments. In addition, we used the Helfrich free energy to model the energetics of membrane deformation, and accounted for the work done against turgor pressure due to changes in volume (Supplemental Text 1). These calculations indicate that the total energy is minimized when filaments orient along the direction of maximal curvature (Fig. 4F) and that, importantly, the energy penalty for incorrectly-oriented filaments is much greater than the energy of thermal fluctuations. Interestingly, this modeling indicates a decrease in energetic preference for the preferred filament orientation as the radius of the cell is increased (Fig. 4F), a prediction in qualitative agreement with our observations of alignment in protoplasts. Furthermore, our calculations indicate that orientation is robust over a large, biologically relevant range of parameters, including the membrane binding energy, filament length, and filament Young’s modulus (Fig. 4G).

These calculations predict that filaments should orient circumferentially both if the membrane deforms to the filament (at low turgor pressures or if filaments are stiff) (Salje et al., 2011), or if filaments deform to the membrane (at high turgor pressures or if filaments are flexible) (Fig. 4E). Our experimental data demonstrates MreB filament alignment across a range of pressures: high within cells, low to none within liposomes, and a pressure between the two within osmotically-stabilized protoplasts. In the absence of turgor pressure, MreB filaments deform liposomes since it is energetically more favorable to bend the membranes than to bend the filaments, as observed in our *in vitro* data (Fig. 4A, S4). However, in live cells, our modeling predicts that MreB filaments cannot deform the inner membrane due to the large turgor pressure, and instead deform to match the greatest principal membrane curvature. Hence filaments create curvature in liposomes and sense it in cells.

### Rod-Shape is Lost in a Global Manner, but Reforms Locally

Together, the above data demonstrate that MreB filaments are sufficient to preferentially orient along the direction of greatest principal membrane curvature. In rod-shaped cells, this direction is along the rod circumference. As filaments move along their length, their orientation constrains the spatial activity of the PG synthetic enzymes such that new cell wall is inserted in a mostly circumferential direction (Hayhurst et al., 2008) to reinforce rod shape (Chang and Huang, 2014; Yao et al., 1999). While the ability of MreB filaments to orient in pre-existing rods can help explain how rod shape is maintained, we also wanted to understand how MreB filaments facilitate the *de novo* formation of rod shape. To explore this, we observed how cells interconvert between spheres and rods.

We first examined how rod shape fails, by growing our TagO-inducible strain at induction levels that produced rods and then reducing the Mg^2+^ concentration to induce them to convert to spheres. This transition revealed that rods convert into round cells by continuous swelling: once a rod begins to widen, it continues to do so until reaching a fully spherical state with no reversion during the process (Fig. 5A). Similar rod to sphere transitions could be attained by holding Mg^2+^ constant while reducing TagO expression. Likewise, cells grown at intermediate TagO induction levels (8-12mM) grew as steady state populations of interconnected rods and spheres, indicating that cells underwent repeated cycles of rod shape formation followed by reversion to spheres (Fig. 1D, E). These results indicate that rod shape can be maintained only as long as the cell wall is sufficiently rigid to resist the internal turgor pressure.

**Figure 5.**
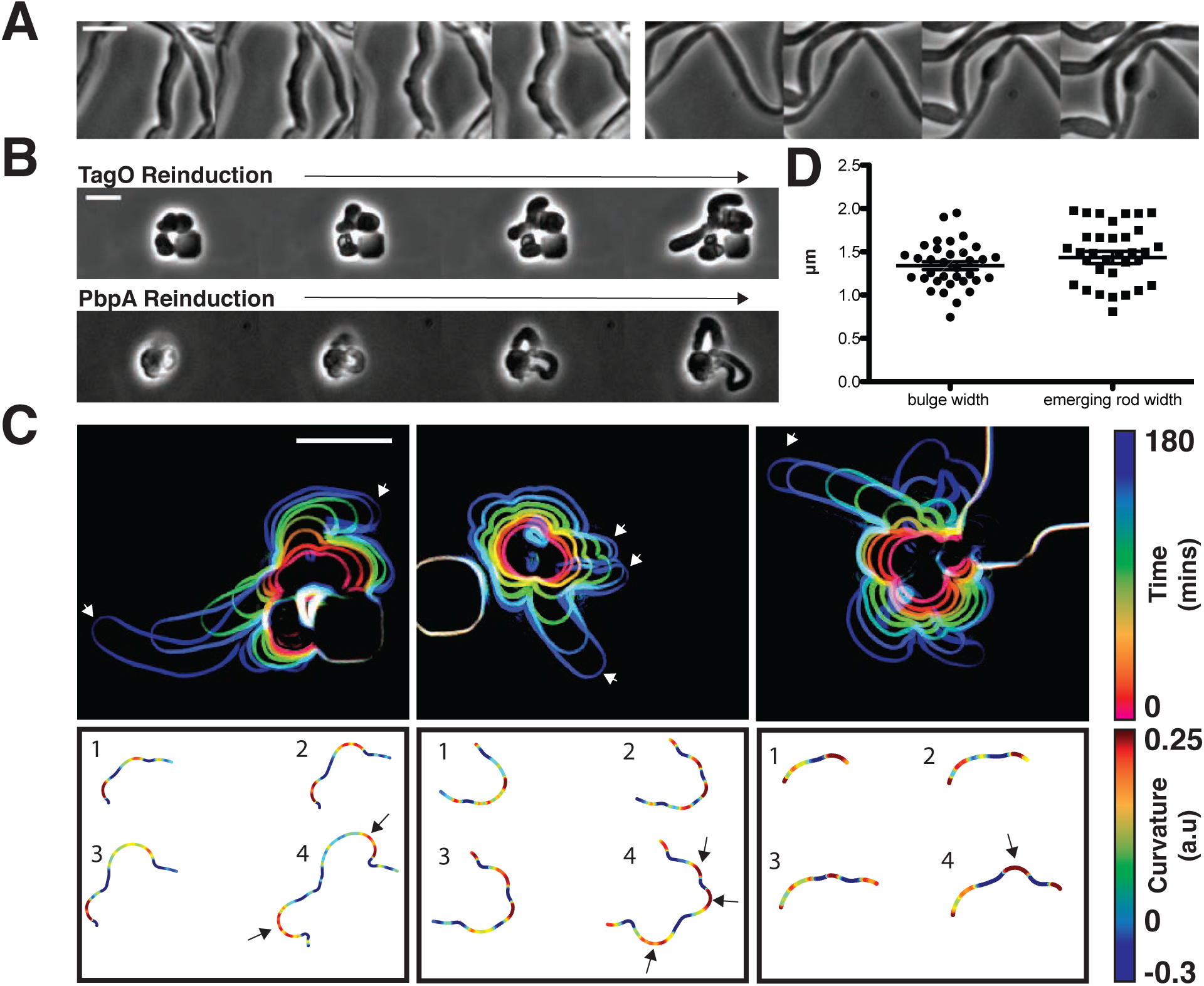
Sphere to rod transitions occur locally and lead to faster growth. **(A)** Loss of rod shape proceeds continuously and without reversals, as shown by BEG300 cells grown in 12 mM xylose, shifted from 1mM Mg^2+^ to 100 μM Mg^2+^ on a pad. Frames are 5 min apart. **(B)** Increases in expression of *tagO* or *pbpA* from depleted spherical cells causes cells to emit rapidly elongating rods from discrete points. (*Top*) BEG300 cells in 20 mM Mg^2+^ were grown in 0 mM xylose for 4h, then transferred to a microfluidic chamber and grown in 0 mM xylose and 20 mM Mg^2+^ for 1h. Following this, *tagO* expression was induced with 30 mM xylose at the first frame. (*Bottom*) BRB785 cells in 20 mM Mg^2+^ were depleted of Pbp2a by growth in 0 mM IPTG for 4h. At the start of the frames, they were transferred to an agar pad containing 1mM IPTG to induce *pbpA* expression. Frames are 30 min apart. **(C)** Plots of cell contours as cells recover from TagO depletion: (*top*) cell outlines are colored in time red to blue (0-180min). White arrows indicate emerging rods; (*bottom*) heat maps of curvature show that rods emerge from small outward bulges (red) flanked by inward curvatures (blue). Black arrows indicate points where emerging rods form. **(D)** The width of initial bulges and the rods that emerge from them are highly similar, indicating the initial deformations may set the starting width of the rods. Error bars are SEM. All scale bars are 5 μm.

We next examined how rod shape forms from round cells. As the recovery of protoplasted *B. subtilis* is so infrequent that it has never been directly visualized (Mercier et al., 2013), we assayed how round cells with preexisting cell walls convert back into rods, using three systems: 1) re-inducing WTA expression within TagO-depleted, spherical cells, 2) holding TagO expression beneath the rod/sphere transition and increasing Mg^2+^ levels, and 3) re-inducing Pbp2a expression in spherical, Pbp2a-depleted cells. In all three cases, rods reformed in a discrete, local manner; spheres did not form into rods by progressively shrinking along one axis, but rather, rods abruptly emerged from one point on the cell, growing more rapidly than the parent sphere (Fig. 5B, Movie S6, Movie S8). This morphology is similar to the initial outgrowth of germinating *B. subtilis* spores (Pandey et al., 2013). We occasionally observed another mode of recovery, occurring when round cells were constrained, or divided into, ovoid or near-rod shapes. Once these near-rod shaped cells formed, they immediately began rapid, rod-like elongation along their long axis (Fig. S5D).

### Rods Form from Local Outward Bulges and Grow Faster than Non-Rod Shaped Cells

We focused on two salient features of the rod shape recoveries: 1) rod shape forms locally, most often at one point on the cell surface, and 2) once a rod-like region is formed, it appears self-reinforcing, both propagating rod shape and growing faster than adjacent or attached non-rod shaped cells.

We first wanted to understand how rod shape initiates *de novo* from spherical cell surfaces. By examining the initial time points of recoveries, we found that rods begin as small outward bulges: local regions of outward (positive Gaussian) curvature flanked by regions of inward (negative Gaussian) curvature (Fig. 5C). These initial outward bulges showed a width distribution similar to that of the later emerging rods (Fig. 5D). Onc these bulges formed, they immediately began rapid elongation into nascent rods, which would then thin down to wild type width over time. Bulge formation and rod recovery were independent of cell division, as cells depleted of FtsZ still recovered rod shape (Fig. S5B). Rather, these bulges appeared to arise randomly, evidenced by the fact that different cells produced rods at different times during WTA or Pbp2a repletion. We conclude that the appearance of a local outward bulge can act as the nucleating event of rod shape formation.

As emerging rods appeared to grow faster than adjacent round cells, we tested if the doubling times of rod-shaped cells were faster than those of non-rods by measuring the doubling times in our inducible TagO strain at different induction levels using both OD_600_ measurements and single cell microscopy under steady state conditions (Fig. 6B, S3C). This revealed a sharp transition in doubling time that matched the conditions of the rod/sphere transition: growth is slow when cells are spheres, yet greatly increases when cells are rods (Fig. S3C, S5A). Furthermore, the doubling times of recovering rods was similar to that of rods at steady state (Fig. S3C).

**Figure 6.**
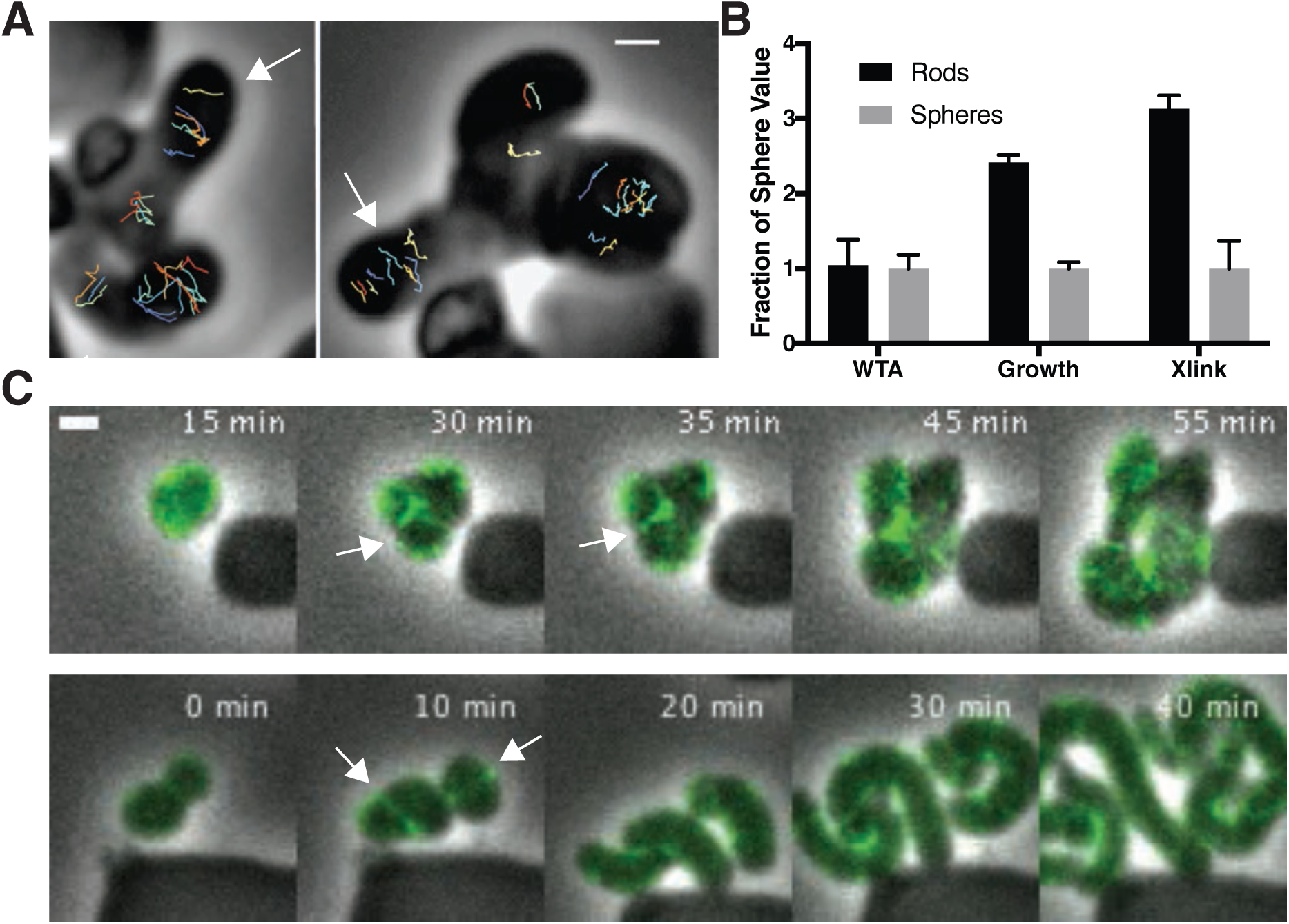
Cell wall crossslinking and growth are coupled to MreB-guided formation of rod shape. **(A)** (*left*) GFP-MreB trajectories during a sphere to rod transition. Emerging rods exhibit oriented MreB motion (white arrows) while attached round cells have unoriented motion. Scale bar is 1 μm. **(B)** Fold change in the teichoic acid incorporation, doubling times (assayed by OD_600_ measurements), and % crosslinked muropeptides of rods (inducible TagO with 30mM xylose in LB with 20 mM Mg^2+^) compared to spheres (grown in LB with 20 mM Mg^2+^). Error bars are SD. See also Figure S3C and S6. **(C)** During shape recoveries, immediately before rod emergence, MreB transiently accumulates in a bright ring where the bulge connects to the parent sphere. See also Figure S5E. Scale bar is 2 μm.

We believe the lower doubling time of rods is likely due to cell shape and not another effect, such as the lack of WTAs, as 1) the doubling time of TagO-depleted cells confined in the microfluidic chambers matched that of wild type cells; and 2) both the doubling times and the boundary of the rod/sphere transition could be equivalently shifted by changing the Mg^2+^ concentration (Fig. 1E, S1, S5A). Combined, these results indicate that rod shape creates local, self-reinforcing regions that are poised for more rapid growth; once any small region of the cell approximates a rod shape, growth of the rod-like region is amplified, growing faster than other regions, and thereby outcompeting non-rod growth at the population level.

### Rod-shape Formation Correlates with Aligned MreB Motion and Increased Glycan Crosslinking

We next sought to determine what features distinguished rods from round cells. As the elongation of rod-shaped cells requires a sufficiently rigid cell wall (Fig. 1D-E, 5A), the self-reinforcing growth of rods could arise from a few mutually compatible sources relative to round cells in our strain: 1) The arrangement of PG strands could be such to reinforce the rod (Amir and Nelson, 2012; Chang and Huang, 2014), 2) WTAs could be preferentially incorporated into rods, or 3) The extent of crosslinking of newly inserted material in the cell wall could be increased so as to make it more rigid (Loskill et al., 2014).

To assay the orientation of newly inserted cell wall, we imaged the motions of MreB as we induced TagO-depleted cells to recover into rods. This revealed that oriented MreB motion correlates with local shape: emerging rods displayed oriented MreB motion even at the initial points of their formation, while attached round parent cells displayed unaligned motion (Fig. 6A, Movie S7). This demonstrates that oriented MreB motion correlates with local geometry and does not arise from a global, cell spanning change. We next examined the overall cellular distribution of MreB in recovering cells with confocal microscopy. This revealed that, immediately prior to rod emergence, MreB transiently accumulated in a bright ring oriented perpendicular to the direction of rod emergence, most often occurring at the interface of the bulge and the round cells (Figure 6C, S5E).

The local reinforcement of rod shape in recovering cells could arise from preferential incorporation of the cell wall rigidifying WTAs. As the WTA ligases have been reported to interact with MreB (Kawai et al., 2011), we tested if rod shape correlated with increased WTA accumulation in emerging rods. To test this, we labeled recovering cells with fluorescently labeled lectins that specifically bind to WTAs (Fig. S6A). Following TagO reinduction, WTAs in recovering cells had a disperse, diffuse distribution around the cell (Fig. S6B), equally present in the cell walls of both rods and spheres (Fig. 6B). To test if the WTA ligases move with MreB, we created fluorescent fusions to these proteins at their native locus and examined their dynamics with TIRFM. We were unable to observe any of the circumferential motions expected if the WTA ligases moved with MreB; instead they appeared to be rapidly diffusing on the membrane (Fig. S6C, Movie S9, Supplementary Text 2).

Next, we used muropeptide analysis to examine if there was a difference in the amount of glycan crosslinking between rods and spheres. This revealed that the PG surrounding spheres was significantly less crosslinked than rods (Fig. 6B, S6D). Thus, the cell wall in rods is more crosslinked, and therefore presumably more load bearing (Loskill et al., 2014). This result fits with the finding that WTA-depleted cell walls are thicker and more irregular (D’Elia et al., 2006). Similarly, studies of plant cell walls have shown that decreased crosslinking makes the cell wall more permeable to water, resulting in swollen, less rigid cell walls (Redgwell et al., 1997)(Ishii et al., 2001).

Previous studies have shown that the rate of PG incorporation is unchanged during the initial phases of depletion of components within the WTA pathway (Pooley et al., 1993). To observe whether both spheres and rods inserted new PG during the process of rod shape recovery in our assay, we used fluorescent D-amino-acids (FDAAs), which crosslink into newly inserted cell wall. We grew TagO-depleted cells in a microfluidic device in the presence of HADA, then switched the media to contain Cy3B-ADA as we re-induced TagO expression. During rod emergence, the old cell wall signal (HADA) remained in the sphere, while the emerging rod was almost entirely composed of new (Cy3B-ADA) material, confirming the discrete nature of rod shape recovery. However, the attached spheres also incorporated Cy3B-ADA, indicating PG synthesis occurs in both rods and spheres during recovery (Fig. S5C).

In summary, this data gives new insights into what properties of the cell wall can be modulated to create and stabilize rod shape: rod shape is not formed by preferential localization of teichoic acids to rods, and both spheres and rods incorporate PG before and during rod shape recovery, in line with reports that PG synthesis is unchanged by the inhibition of WTA synthesis (Pooley et al., 1993). Rather, the only differences between rod shaped and round cells we observed were 1) oriented motion of MreB in rods, coupled with 2) an increased crosslinking of the inserted glycans. Thus, it appears that not only does MreB direct glycan insertion into circumferential hoops, but also these strands are more crosslinked, both properties are expected to increase the strength of the rod sidewalls (Loskill et al., 2014)(Yao et al., 1999). It may be that these two attributes are mechanistically linked, and a more oriented arrangement of glycan strands might provide a more optimal arrangement of peptides for crosslinking reactions. We note that as PG and WTA precursors share a common lipid carrier and WTAs affect hydrolase activity (Kasahara et al., 2016), their depletion may cause other rod-shape inhibiting PG abnormalities that we cannot observe.

## Discussion

The above experiments give new insights into the mechanism by which MreB builds rod shape. First, the curved ultrastructure of MreB filaments causes them to orient and move along the direction of greatest membrane curvature, inserting material in that direction. Second, both the formation and propagation of rod shape occurs by a local, self-reinforcing process: once a local region of rod shape forms, it propagates more rod shape. Finally, as far as we can determine, the primary differences between the growth of rods and non-rods is the circumferential orientation of MreB motion and increased glycan crosslinking.

Combined, these findings indicate that MreB filaments function as curvature-sensing rudders, a property that allows them to organize cell wall synthesis so that it builds rod shape: MreB filaments orient along the greatest membrane principal curvature, thereby constraining the activity of the associated PG synthases so that, as they move via their synthetic activity, they deposit highly crosslinked glycans oriented in the direction of that curvature, and this arrangement of glycan insertion reinforces rod shape. Even during the initial stages of rod shape formation, oriented MreB motion and rod shape always coincide, and the intrinsic curvature of MreB filaments suggests these properties cannot be uncoupled. This coupling appears to be an essential component of the Rod system: by linking filaments that orient along the greatest principal curvature to cell wall synthetic enzymes reinforcing that curvature, the Rod complex creates a local, self-organizing system that allows bacteria to both maintain rod shape and also establish rod shape *de novo*.

In established rods, we propose that MreB maintains and propagates rod shape via feedback between existing shape, filament orientation, and subsequent shape-reinforcing PG synthesis. As rod-shaped cells grow (Fig. 7A1), MreB filaments orient along the more curved axis around the bacterial width (Fig. 7A2). Because MreB filaments always translocate along their length (Olshausen et al., 2013), filament orientation constrains the activity of the associated PG synthases such that new cell wall is inserted in bands predominantly oriented around the width of the rod (Fig. 7A3). This circumferential insertion of glycan strands, combined with a high level of crosslinking between them, yields a highly connected, anisotropic arrangement of material that reinforces rod shape (Fig. 7A1), which allows continued MreB filament orientation. This feedback loop can continue as long as the material within the rod sidewalls is sufficiently rigid to withstand the stresses arising from the internal turgor pressure, allowing the rod shape to be robust once it is formed.

**Figure 7.**
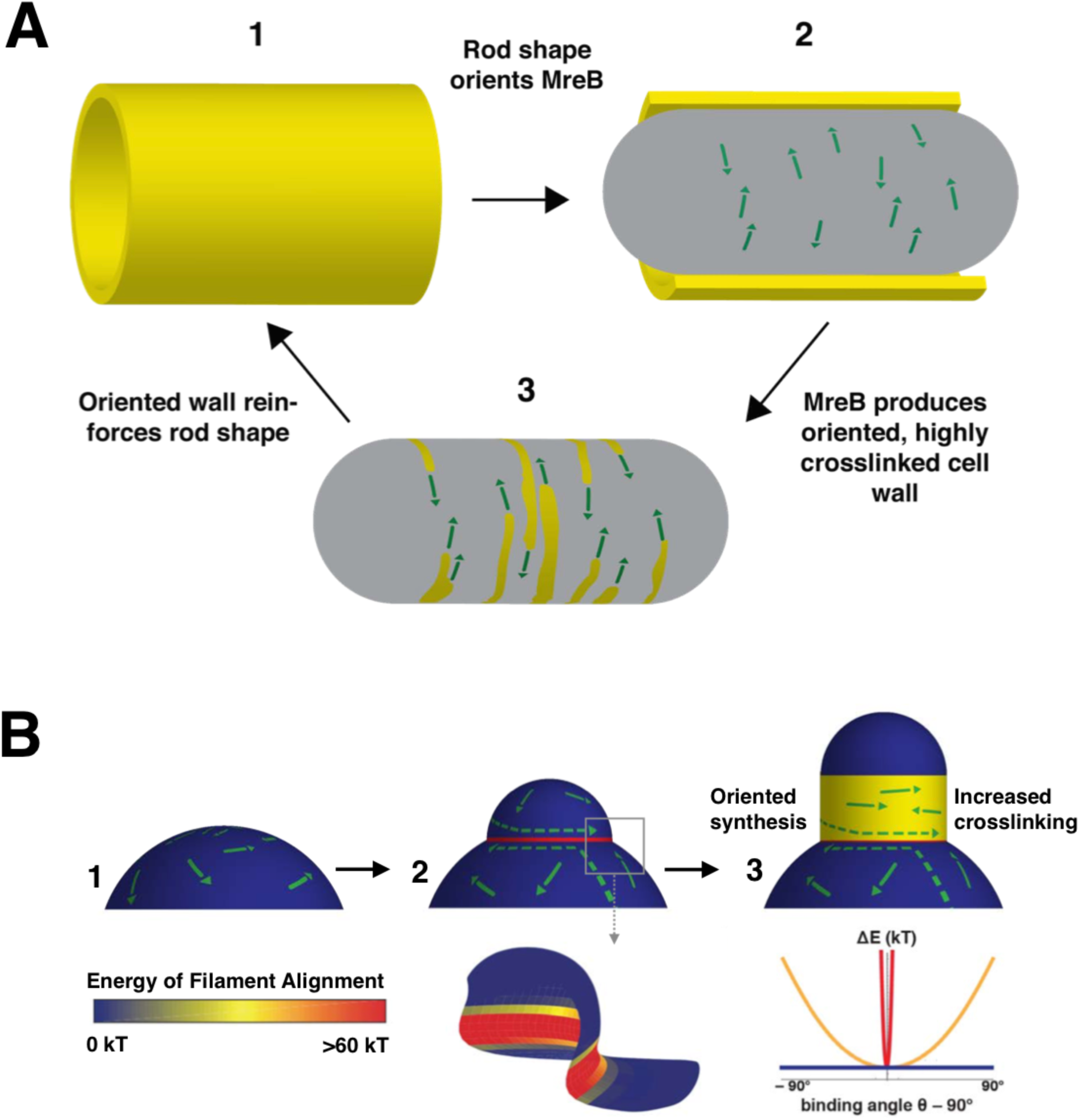
Model for how MreB filament orientation along the greatest curvature both maintains and establishes rod shape. **(A)** Rod-shaped cells present a single curved axis along which MreB filaments orient (1). This orientation determines the direction of MreB motion (2), thus orienting the insertion of new cell wall material around the rod, and allowing an increased crosslinking between strands (3). This highly cross-linked, circumferential arrangement of cell wall material reinforces rod shape (1), leading to more aligned MreB filaments, thus creating a local feedback between the orientation of MreB filaments, oriented cell wall synthesis, structural integrity of the rod, and overall rod shape. **(B)** MreB motion in spherical cells is isotropic (1), but the introduction of an outward bulge (2, upper) creates a curved geometry (red) at the neck of the bulge that initiates rod shape formation. Due to the high energy of alignment in this region, (2 lower and chart), any filaments that encounter the neck of the bulge would prefer to align to point around the neck rather than cross it, creating a ring-shaped region of aligned MreB motion that nucleates rod formation. Repeated rounds of oriented synthesis around the ring could initiate the elongation of a rod from the initial bulge site (3), beyond which rod shaped elongation would be self-sustaining. Colors correspond to the difference of alignment energies along the two principal curvatures at the negatively-curved neck region (red), flat regions with one dimension of curvature (yellow), and the positively-curved sphere/bulge (blue).

The coupling between the local sensing and reinforcement of differences in principal curvature could also allow the *de novo* formation of rod shape. In round cells, there is no difference in principal curvatures (Fig. 7B1), so MreB motion is isotropic. Rods do not form by squeezing these round cells across one axis, rather we observe them forming by the amplification of local rod-like regions. Given the rapid timescale of our recoveries, the Rod system appears poised to propagate any shape variations that create curved regions favorable to oriented MreB motion. Once regions of oriented motion are established, they self-propagate and elongate, creating a new rod shape and thus continued oriented MreB motion. The most common shape variation we observe preceding rod emergence is small outward bulges flanked by regions of inward curvature (Fig. 5C). It remains to be determined how these initial bulges form and what cellular factors are involved in this process. They could arise from local changes in cell wall stiffness: a local softening of the cell wall has been observed to induce the rod shaped outgrowth of germinating fission yeast spores (Bonazzi et al., 2014).

The geometry at the interface of these outward bulges plays a central role in our model of rod shape formation. In three dimensions, the intersection at the bulge and the sphere creates a geometry that can establish a zone of aligned filaments: while both the parent sphere and the outward bulge have principal curvatures in the same direction (positive Gaussian curvature), the intersection of the sphere and bulge creates an interface with strong differences in principal curvatures, one inward, and one outward (negative Gaussian curvature). Upon entering these negatively curved regions it is energetically unfavorable for the inwardly curved MreB filaments to deviate from their preferred binding orientation, as our modeling indicates that this region presents a steep well in the energy profile for alignment (Fig 7B2 and Supplemental Text 1). Thus, filaments moving into this rim from either side would reorient to move along it, creating a concentrated band of filaments moving around the bulge neck. This concentrated ring of oriented MreB filaments may then construct a local region of rod shape that subsequently self-propagates into an emerging rod (Fig. 7B3). In support of this hypothesis, immediately preceding rod shape formation, we observe concentrated bands of MreB transiently appearing at the neck of emerging bulges (Fig. 6C, S5E). Likewise, similar patterns of MreB accumulation at points of negative Gaussian curvatures have been observed in recovering *E. coli* L-forms (Billings et al., 2014).

The common observation of MreB accumulation at the necks of rod-producing bulges in both *E. coli* and *B. subtilis* hints at a solution to an outstanding discrepancy: Why do inwardly curved MreB filaments show an enriched localization at negative Gaussian curvatures (inward dimples or the more curved faces of bent cells) (Billings et al., 2014; Renner et al., 2013; Ursell et al., 2014), and how is this enrichment maintained as filaments move around the cell? The finding that MreB filaments align along the greatest curvature poses a solution: If the sharpness of filament alignment changes in response to the difference in principal curvatures in each region they pass through, areas of negative Gaussian curvature may act as points that focus the subsequent motion of filaments so that, on average, more filaments pass through these regions.

The tendency of MreB to align and move along the direction of greatest principal curvature may also explain the absence of MreB at cell poles. Consistent with our model for binding, we observed MreB filaments bound to the round poles of liposome tubes *in vitro* (Fig. 4A). In the cell, however, MreB filaments move directionally, and filaments entering the symmetrically curved pole in any orientation would quickly translocate out into the cylindrical cell body where they would reorient along the single direction of curvature.

While rod-shaped cells show both an increased rate of growth and oriented MreB motion, it is unlikely these phenomena are mechanistically linked. Rather, the decreased rate of growth of non-rods likely arises from a downstream effect of the lack of rod shape on cell physiology. Indeed, many spatial processes in *B. subtilis*, such as chromosome segregation and division site selection, read out and partition along the long axis established by rod shape (Jain et al., 2012). Thus, the slower doubling times observed in non-rod shaped cells may arise from the improper spatial organization of these processes, or stress responses to this spatial disarray.

As the curvature of membrane-bound MreB filaments (200nm) observed *in vitro* is much greater than the cell diameter (900nm), these findings suggest that the curvature of MreB filaments does not define a specific cell radius; rather filament curvature acts to orient PG synthesis to maintain (Harris et al., 2014) or reduce cell diameter. If the curvature of MreB filaments reflects the smallest possible cell diameter, bacterial width may be specified by opposing actions from the two spatially distinct classes of PG synthases: a decreasing, “thinning” activity from the action of MreB and its associated SEDS family PG synthases, and an increasing “fattening” activity from the non-MreB associated Class A PG synthases.

## Conclusion

To construct regular, micron-spanning shapes made of covalently crosslinked material, nature must devise strategies for coordinating the activities of disperse, nanometer-scale protein complexes. This work reveals that the role of MreB in creating rod shape is to locally sense and subsequently reinforce differences in principal curvatures. The local, short-range feedback between differences in curvature, MreB orientation, and shape-reinforcing cell wall synthesis provides a robust, self-organizing mechanism for the stable maintenance and rapid reestablishment of rod shape, allowing the local activity of short MreB filaments to guide the emergence of a shape many times their size.

## Author Contributions

All authors contributed intellectual and editorial work to the development of the manuscript. S.H., C.N.W., and E.C.G. designed, performed, and interpreted experiments. P.S., T.I., and J.L. contributed the electron microscopy work shown in Fig. 4. F.W. and A.A. contributed biophysical modeling work shown in Fig. 4, Fig. 7, and supplemental text. L.D.R performed microfluidic device fabrication. K.S. and S.W. performed peptidoglycan crosslinking analysis. Y.S. wrote software for growth analysis. A W. B. performed the FtsZ inhibition experiments.

## Acknowledgements

We thank T. Norman, N. Lord, and M. Cabeen for help with PDMS molds and microfluidics; M. Dion and M. Kapoor for strains; E. Kuru, Y. Brun, and M. VanNieuwenhze for FDAAs; G. Billings, S Van Teeffelen, and T. Ursell for helpful discussion; T. Bharat for help with electron microscopy, and S. Layer for advice and inspiration. This work was funded by National Institutes of Health Grant DP2AI117923-01, a Smith Family Award, a Searle Scholar Fellowship (to E. G.), a Sloan Foundation Award (to A.A), an NSF GFRP to (F. W), the Medical Research Council (U105184326 to J.L.) and the Wellcome Trust (095514/Z/11/Z to J. L.). This work was performed in part at the Center for Nanoscale Systems at Harvard University, supported by NSF ECS-0335765. S.H. is a HHMI International Student Research Fellow. K.S. was funded by NIH R01 GM076710.

## Supplemental Figures

**Figure S1.**
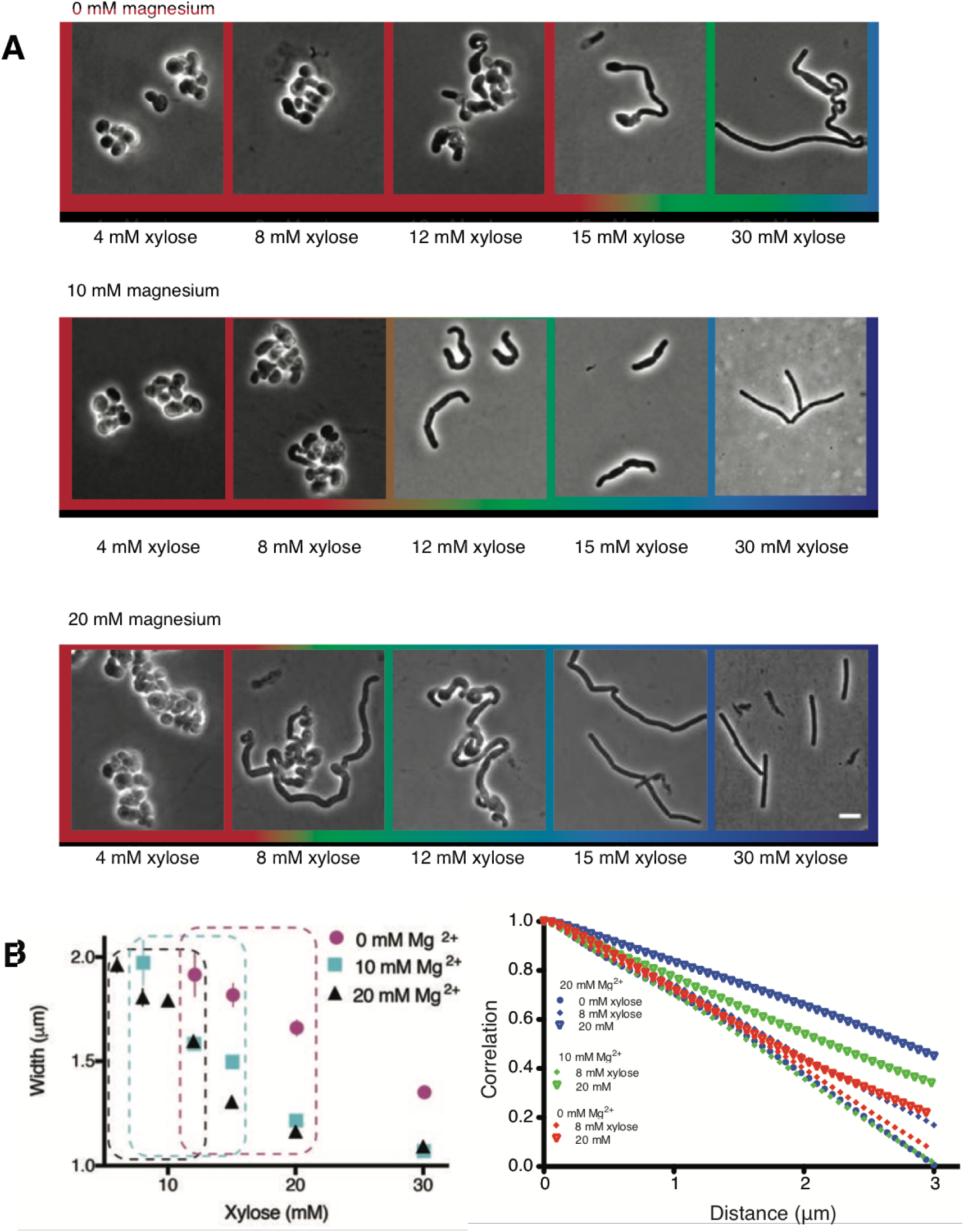
Varying magnesium levels in the growth medium changes cell shape. Related to Figure 1. **(A)** TagO inducible cells grown in LB supplemented with varying Mg^2+^ levels (0, 10 and 20 mM), show similar trends in cell shape across increasing xylose concentrations, with the appearance of more rod-shaped cells that become thinner as xylose levels increase. Exogenous Mg^2+^ reduces the amount of TagO induction needed for rod shape, evidenced by shift in the amount of xylose required to form rods as Mg^2+^ is increased. (Color Outlines: Blue = rods, Green = Mixed rods and non-rods, Red = non-rods). **(B) *Left*** Plot of cell width as a function of TagO induction at different Mg^2+^ concentrations (error bars are SEM). Areas not plotted at lower xylose levels are regions where cells are round, with no width axis. Dotted rectangles mark conditions where both round cells and wide rods exist. Error bars are Standard Error of the Mean (SEM). ***Right*** At low xylose and magnesium levels, tangential correlation along the cell contours falls off faster, indicating loss of rod shape. Correlation of angles was calculated as described in methods. The curves shown are population averages of tangential correlations at selected xylose and magnesium concentrations. A cutoff of 3 μm is applied as this is the mean cell length of *B. subtilis.*

**Figure S2.**
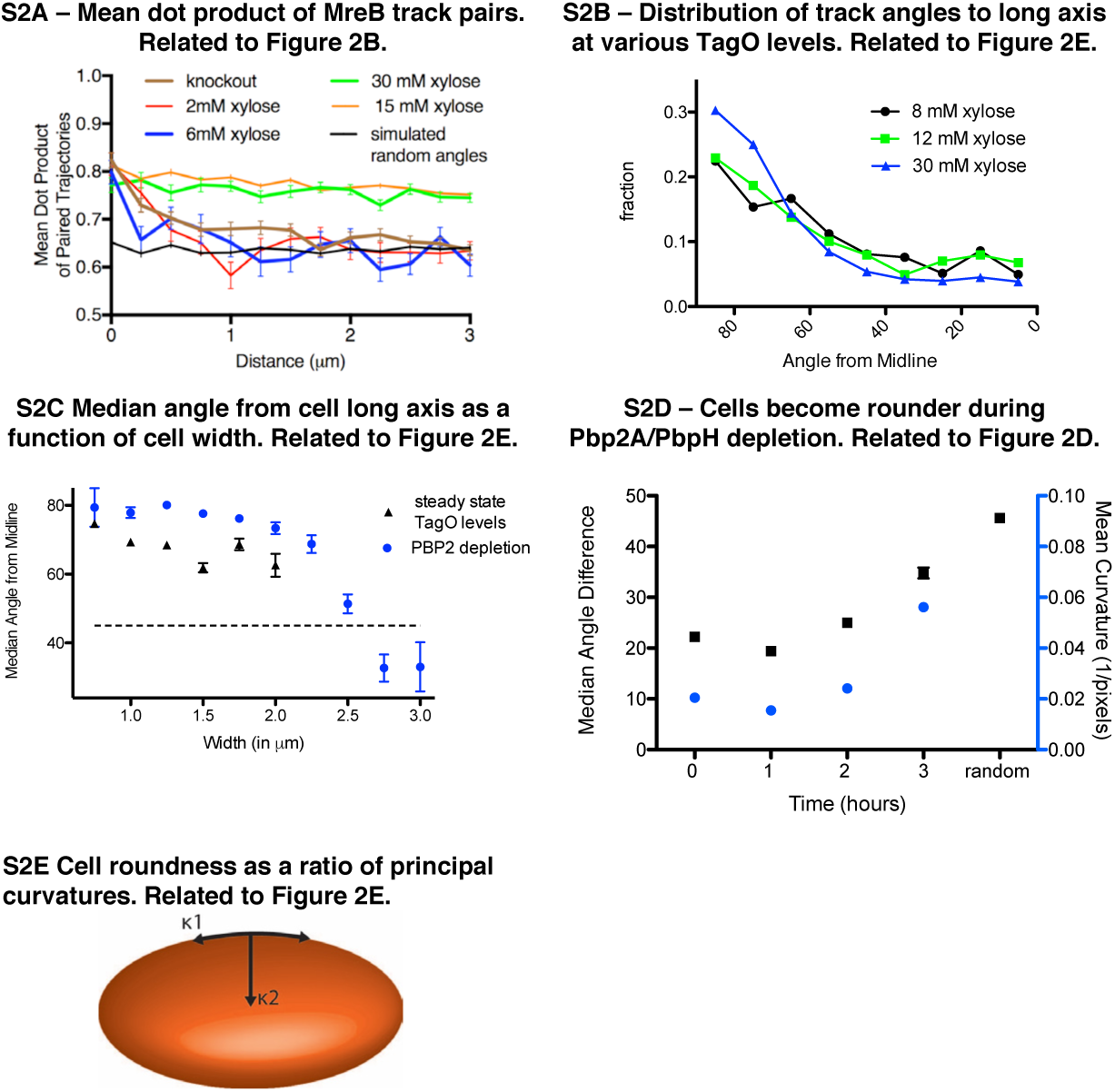
(A-E) – Relationships between cell width, MreB orientation, and cell shape. Related to Figure 2. **(A)** The mean dot product of MreB track pairs vs. distance between the pairs was calculated and binned at 0.25 μm intervals. This shows a high alignment between pairs across the cell length for rods at high xylose (15 and 30mM). Round cells (2 and 6mM xylose) show a high alignment at very short distances (< 500 nm), beyond which alignment falls off rapidly, approaching the value expected for randomly oriented angles (black line represents a simulation of a uniform angular distribution). **(B)** MreB filament motion is predominantly circumferentially oriented over a range of xylose levels (8-30mM) even though cells show varying widths. At 8 and 12 mM xylose, cells are a mix of rods and spheres, and therefore angles were only calculated for cells with identifiable long axes. **(C)** Median angle from the long axis of cells as a function of cell width at steady state TagO levels and PBP2a depletions (shown separately). **(D)** Mean sidewall curvature of cells increases during a Pbp2a/PbpH depletion (blue circles), along with a decrease in aligned MreB motion (black squares). **(E)** The principal curvatures along the cell length (κ1) and cell width (κ2) are calculated and the ratio κ1/κ2 is taken as a measure of cell roundness. Cells become round as this ratio approaches 1. All error bars are SEM.

**Figure S3.**
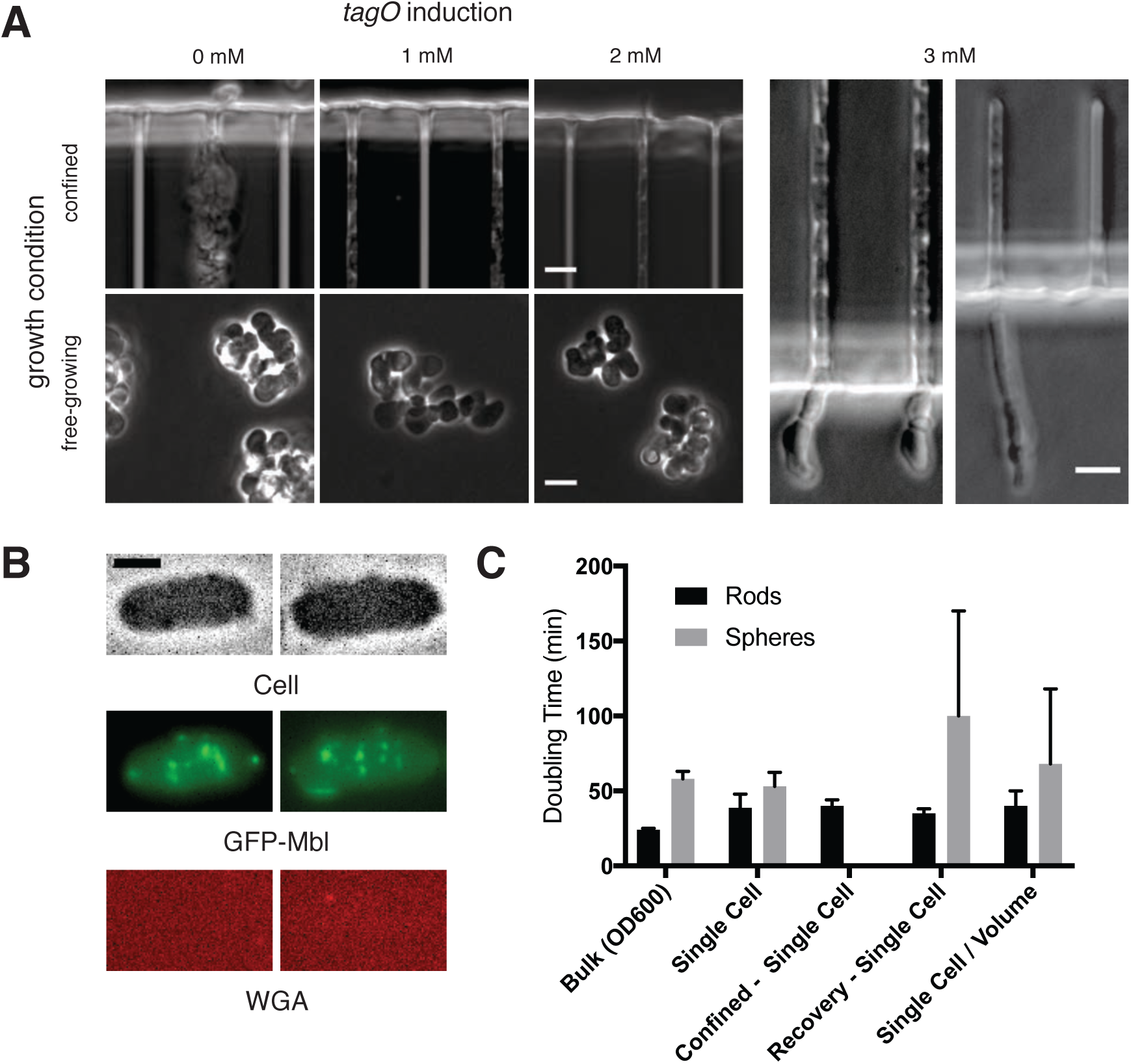
TagO Levels and Confinement. Related to Figure 3. **(A)** Microfluidic confinement controls cell shape in cells with low TagO levels. (*Left*) Phase contrast images of BEG300 grown under differing teichoic acid induction levels in bulk culture or confined in chambers. *(Right)* Cells swell upon escaping from confinement. Swelling is visible both at initial stages of depletion, corresponding to when MreB movies were collected (left panel, cf. Figure 3C and Supplementary Movie 3), or at longer stages when cells were chained (right panel). Scale bars = 5 μm.**(B)** Panels showing images of phase contrast, GFP-Mbl, and Alexa455-conjugated WGA of a representative protoplast confined into rod shape. MreB filaments are aligned along the cell circumference, (middle panel), but do not regrow cell wall as indicated by the lack of signal in the Alexa455 channel. Scale bars are 2 μm. **(C)** Doubling time of BEG300 in different conditions. “Bulk” indicates cultures grown in liquid suspension and measured by OD_600_. “Single Cell” indicates cells were grown under agarose pads, with doubling time measured by assaying the change in cell area over time using phase contrast microscopy. “*Confined - Single Cell*” indicates the doubling time of cell area of TagO-depleted cells confined into rod shape in microchambers as in Figure 3A and S3A; “*Recovery – Single Cell*” is the single-cell doubling time (in volume) of TagO-depleted cells during rod shape recovery in a cellASIC microfluidic device as in Figure 5B. Note that spherical cells in these recoveries show a slower doubling time with a larger standard deviation due to a subpopulation of cells dying during the experiment; “*Single cell / Volume*” indicates the doubling time of the volume of single cells grown in a cellASIC microfluidic device. As this chamber has a fixed Z height, cell volume can be approximated from measures of the 2D area. Error bars are standard deviation.

**Figure S4.**
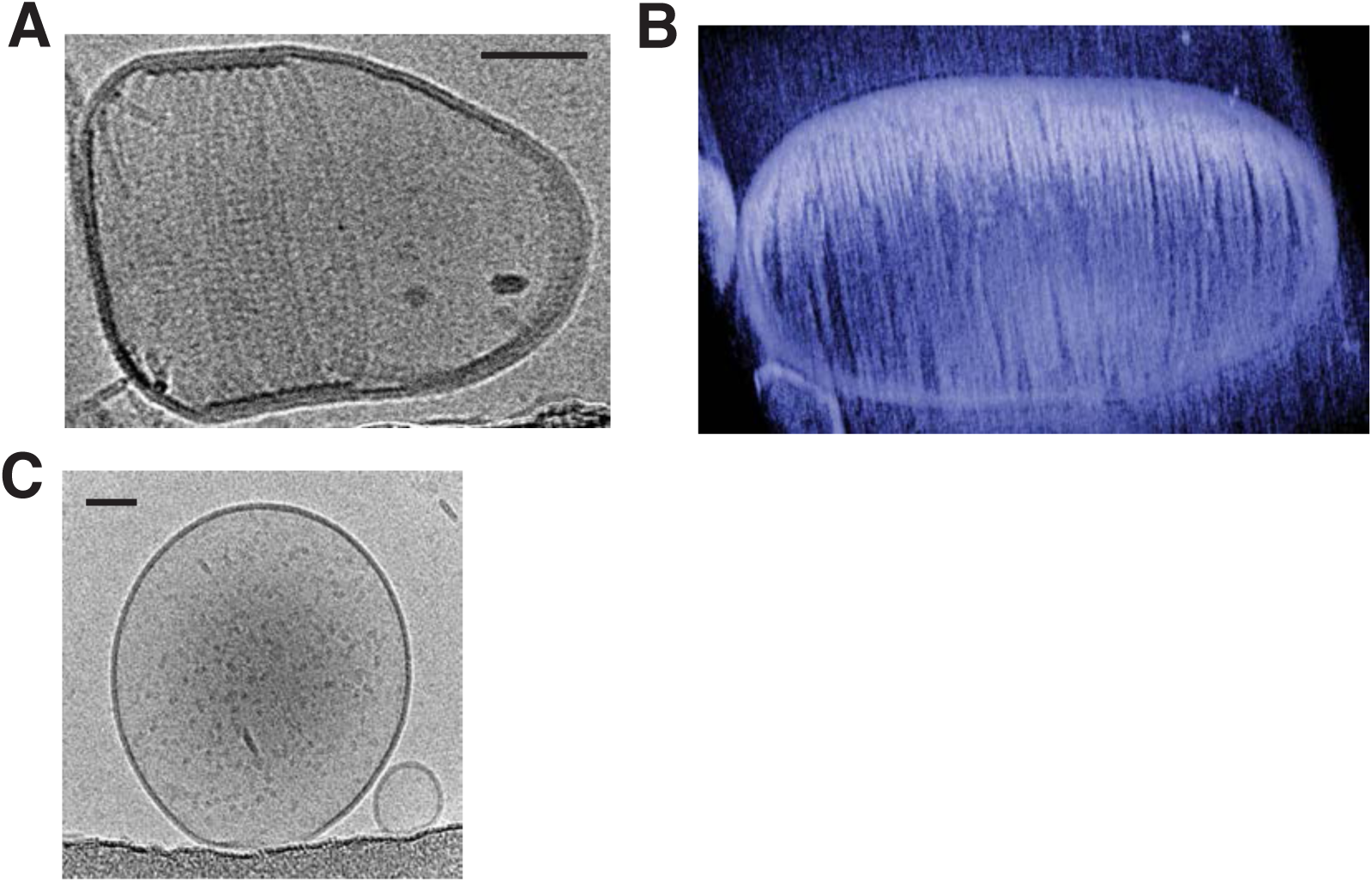
*T. maritima* MreB filaments assembled in liposomes support alignment to rod axis. Related to Figure 4. **(A)** The arrangement of MreB filaments inside the liposome is helical. Scale bar is 50 nm. **(B)** Many long *maritima* MreB filaments inside an artificial liposome, assembled *in vitro* and imaged by electron tomography. Almost the entire inner surface of the liposome is covered with filaments, leading to deformation of the normally spherical liposome. Corresponding movie: SM5, first part. Scale bar is 50 nm. **(C)** Control showing that liposomes are spherical in the absence of MreB. Scale bar is 50 nm.

**Figure S5.**
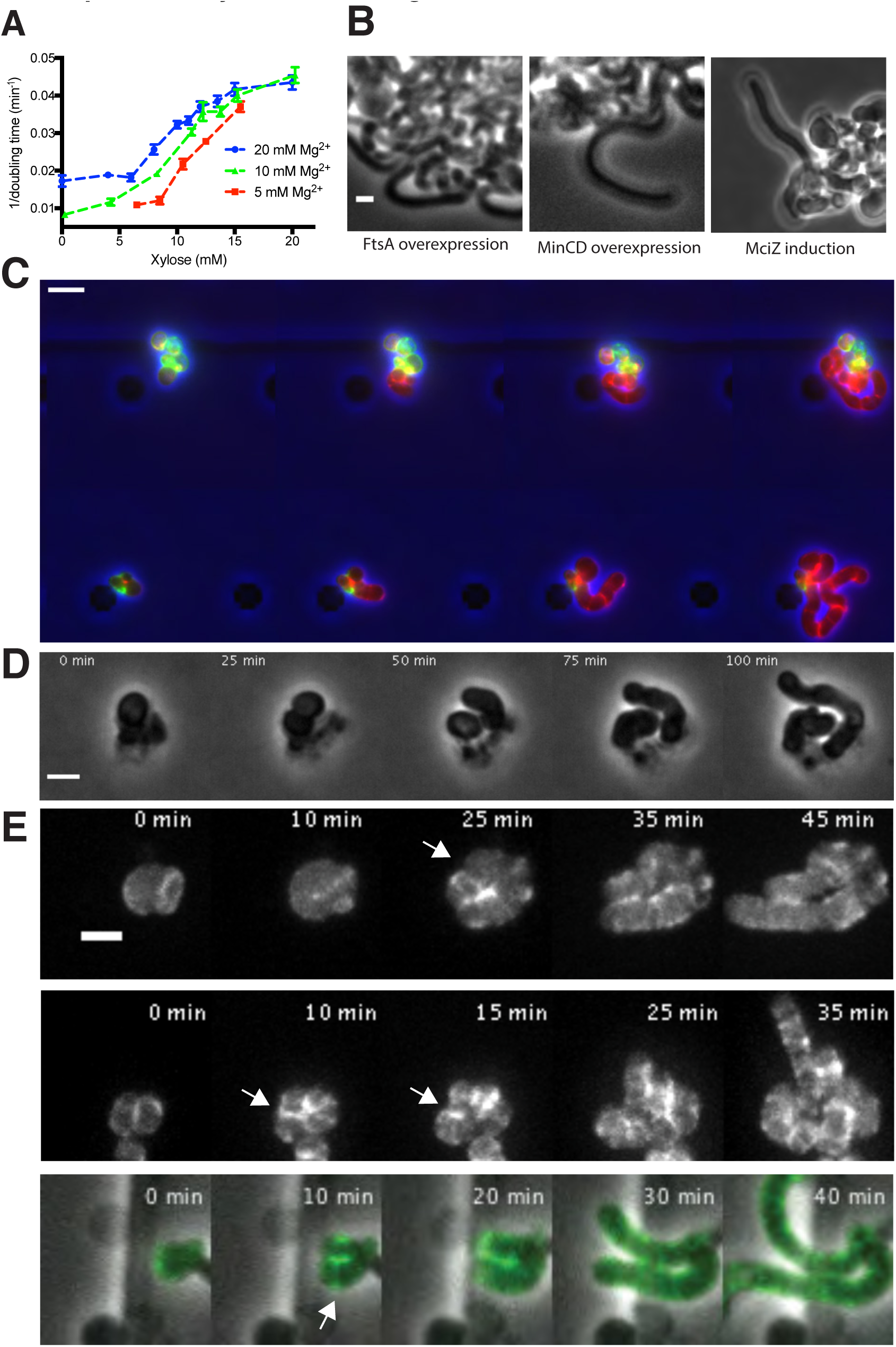
Growth of rod-shaped and spherical cells measured by doubling times, and rod shape recovery. Related to Figure 6. **A)** Rate of doubling (1/doubling time), calculated from OD_600_, increases with increasing levels of TagO as round cells become more rod-like. Increasing Mg^2+^ causes these curves to shift leftward, as Mg^2+^ stabilizes rod shape in combination with WTAs (see Fig S1A). Error bars are SEM. **(B)** Rod shape recovery occurs in the absence of cell division and FtsZ filaments. Inhibition of FtsZ filaments was conducted by three means: FtsA overexpression (bAB388, grown with 1 mM IPTG and 60 mM xylose), MinCD overexpression (bAB327 grown with 1 mM IPTG and 60 mM xylose), and MciZ induction (bAB343 grown in 1 mM IPTG and 30 mM xylose). In all cases, cells recovered rod shape. **(C)** Pulse chase labeling with FDAAs during TagO recoveries indicates that while emerging rods are composed of new cell wall, both spheres and rods incorporate new cell wall material. BCW82 was grown in a microfluidic chamber with 0 mM xylose, 20 mM Mg^2+,^ and 3 μM HADA (green). Prior to imaging the medium was switched to 30 mM xylose (to induce TagO expression), 20 mM Mg^2+^ and 3 μM cy3B-ADA (red, to visualize new cell wall incorporation). Cell outline (from phase) is shown in blue. Scale bar is 5 μm; frames 30 min apart. **(D)** A montage of a rod shape recovery occuring after a division that produced an ovoid, near rod-shaped cell that subsequently elongated as a rod. This example taken from an experiment with BRB785, where Pbp2a was first depleted (in a *pbpH* null) to make round cells, then Pbp2a was reinduced with 1 mM IPTG. See also bottom panel of S5E. **(E)** MreB localizes in a ring-like structure (white arrows) at the neck of emerging bulges, immediately prior to rod shape formation. BEG300, containing GFP-MreB was depleted of TagO by growing in bulk culture in media lacking xylose. Cells were then loaded into a cellASIC device, and grown for 2 hours in the same media with 1 mM IPTG added to induce GFP-MreB expression. At the start of imaging, the media was switched to 30 mM xylose to induce TagO expression, and Z-stacks of GFP MreB were taken using a spinning disk confocal every 5 minutes. Shown is the maximal intensity projection of entire cell.

**Figure S6.**
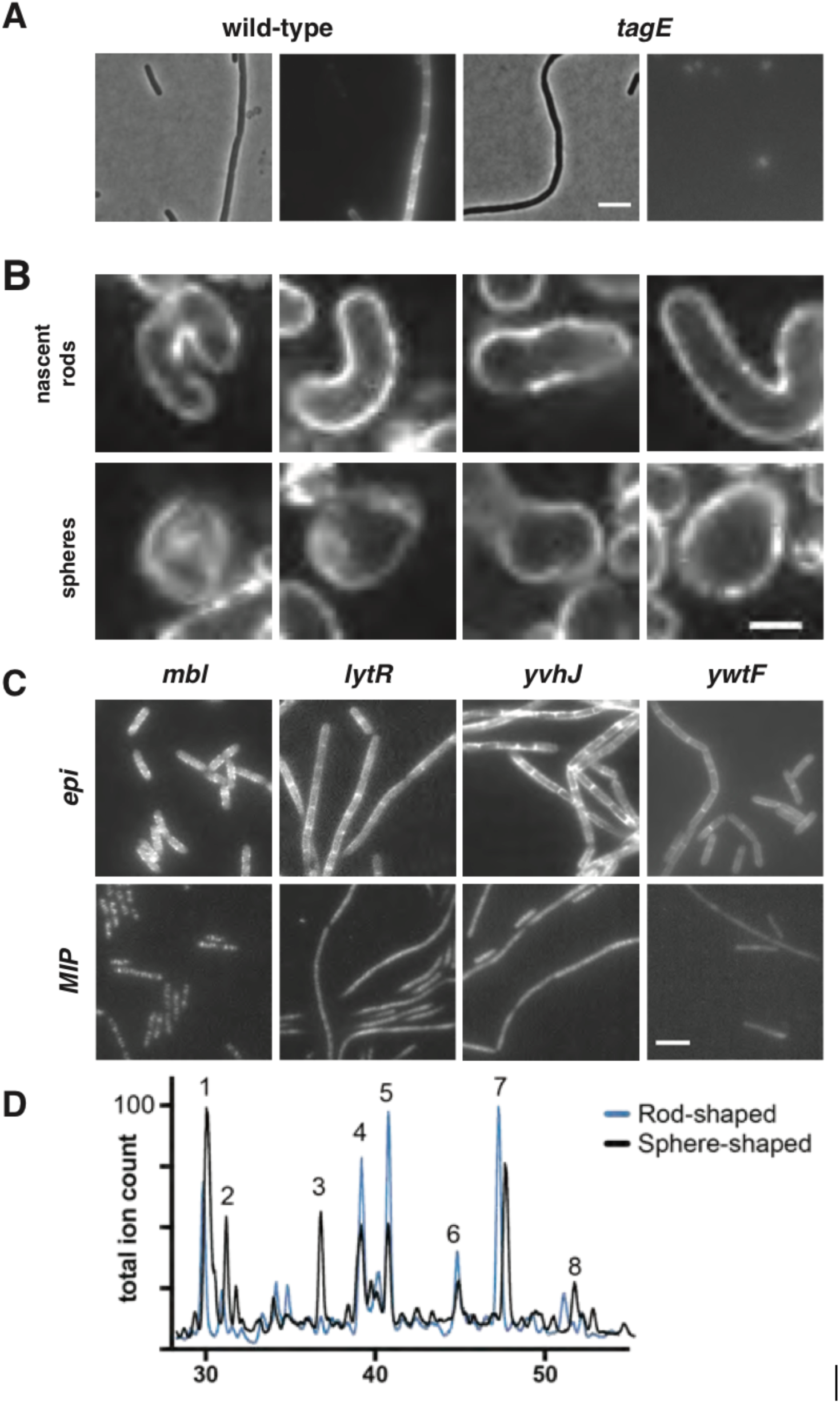
ConA staining of PY79δtagE and muropeptide analysis of BEG300. Related to Figure 6B. **(A)** Concanavalin A conjugated to Alexa Fluor 647 specifically stains wall teichoic acids and localizes uniformly around the entire cell during sphere to rod transitions. A comparison of PY79 and δ*tagE* cells stained with ConA-A647 reveals specificity of ConA for WTAs. The *tagE* gene is responsible for the glycosylation of wall-teichoic acids, rendering them susceptible to ConA binding. Scale bar is 5 μm. **(B)** Fluorescence intensity of Alexa647-ConA, is comparable in rod shaped and round cells during sphere to rod recoveries. Furthermore, the WTA incorporation is diffuse, and not banded or localized. BEG300 was grown in bulk culture and rod shape was induced by increasing TagO levels (0 to 30 mM xylose). Scale bar is 2.5 μm. **(C)** Wall teichoic acid ligases diffuse homogenously across the cell surface. Ligases were tagged at the native locus and promoter with msfGFP and imaged continuously with 100 ms exposures. Epifluorescent images were collected under oblique laser illumination and maximal intensity projections (MIPs) were created over 100 frames of TIRF illumination (see Movie S8). For comparison, msfGFP-tagged Mbl, which localizes in discrete patches, is shown. Scale bar is 5 μm. **(D)** Overlaid LC/MS traces corresponding to reduced muropeptides isolated from the sacculi of rod-shaped and sphere-shaped *B. subtilis*. The total ion count was scaled to the highest peak for each trace, and muropeptides were identified under the total ion count. Peaks 1-3 correspond to uncrosslinked muropeptides; peaks 4-8 correspond to crosslinked muropeptides. Muropeptides identified included the following: a tripeptide (GlcNAc-MurNAc-L-Ala-D-Glu-m-DAP by its [M+1]=871.3 (Exact mass=870.3) were identified under peak 1; a dipeptide (GlcNAc-MurNAc-L-Ala-D-Glu) was identified under peak 2 [M+1]=699.7 (Exact mass=698.7); a tetrapeptide (GlcNAc-MurNAc-L-Ala-D-Glu-m-DAP-D-ala by its [M+1]=941.2 (Exact mass=940.2) was identified under peak 3; crosslinked, dimeric muropeptides were identified in peaks 4-7 containing the tetrasaccharide (GlcNAc-MurNA-L-Ala-D-Glu-m-DAP-D-ala-m-DAP-D-Glu-L-Ala_MurNAc-GlcNAc) by [M+1]=1795.8 and [M+2]/2=898.4; (Exact masses=1794.8); and peak 8 corresponded to a crosslinked, trimeric muropeptide containing two tetrapeptides (GlcNAc-MurNA-L-Ala-D-Glu-m-DAP-D-ala) and a tripeptide (GlcNAc-MurNAc-L-Ala-D-Glu-m-DAP); [M+3]/3= 906.9 (Exact mass=2717.7). To calculate the amount of each species, peaks were integrated by their extracted ion chromatograms. Crosslinked species were calculated using previous literature quantification methods (Glauner et al., 1988). In the rod-shaped *B. subtilis*, 41% crosslinking was observed; in sphere-shaped *Bacillus subtilis*, only 11% crosslinking was observed.

## Materials and Methods

### Overnight culture growth

All *B. subtilis* strains were prepared for experimentation as follows: strains were streaked from −80°C freezer stocks onto lysogeny broth (LB) agar plates. Following >12 hours of growth at 37°C, single colonies were transferred to serially diluted overnight bulk liquid cultures in LB supplemented with 20 mM magnesium chloride, placed on a roller drum agitating at 60 rpm, and grown at 25°C. After >12 hours growth to OD_600_ < 0.6, these starter cultures were transferred to or inoculated into subsequent growth conditions. All strains with *tagO* under inducible control were grown overnight in the presence of 30 mM xylose unless otherwise noted.

### Single cell and bulk doubling time measurements

For the experiments in Fig. 6B and Fig. S3C, BEG300 cells were inoculated in the indicated medium (LB with 20 mM MgCl_2_ unless otherwise stated) from logarithmic phase overnights; “rods” were grown from a low dilution with 30 mM xylose, and “spheres” were grown with 0 mM xylose.

For bulk culture doubling time measurements, doubling times were calculated from the slope of a graph of time vs. dilution for a succession of serial dilutions of a given strain. Time, the dependent variable, was taken as the time for a given dilution to pass the OD cutoff of OD_600_ = 0.20.

Single cell measurements were made in three ways.

i) Spherical and rod-shaped cells were allowed to grow on agarose pads made with LB supplemented with 20 mM MgCl_2_. 30 mM xylose was added to agarose pads for rod-shaped cells. Cells were imaged every 2 minutes for 4 hours with phase contrast microscopy as described in the section below.

ii) Spherical and rod-shaped cells were grown in the CellASIC B04A plate in LB supplemented with 20 mM MgCl_2_ for spherical cells and LB supplemented with 20 mM MgCl_2_ and 30 mM xylose for rod-shaped cells. The CellASIC unit confined the cells in the Z dimension due to the fixed height of the ceiling. Cells were imaged every 10 minutes for 2 hours using phase contrast microscopy as described in the section below.

iii) For cells growing in the mother machine microfluidic device (see below), the expansion of the cell length along the channel was quantified using FIJI (Schindelin et al., 2012; Schneider et al., 2012); only the cells closest to the mouth of the channel were counted. Since cells were always oriented along the length of the channel (see Fig. 3A, S3A), changes in expansion in this dimension accounted for all growth.

### Imaging – phase contrast microscopy

Phase contrast images were collected on a Nikon (Tokyo, Japan) Ti microscope equipped with a 6.5 μm-pixel Hamamatsu (Hamamatsu City, Japan) CMOS camera and a Nikon 100X NA 1.45 objective. Cells were collected by centrifugation at 6,000 x *g* for 2 min and re-suspended in the original growth medium. Unless otherwise specified, cells were then placed on No. 1.5 cover glass, 24 x 60 mm, under a 1 mm thick agar pad (2-3% agar) containing LB supplemented with 20 mM magnesium chloride. Unless otherwise noted, all cells were imaged at 37°C on a heated stage.

### Imaging – MreB particle tracking

Images were collected on a Nikon TI microscope with a 6.5 μm-pixel CMOS camera and a Nikon 100X NA 1.45 objective. Cells of strain BEG300 were grown overnight in LB supplemented with 30 mM xylose, 20 mM magnesium chloride, 1 μg/mL erythromycin, and 25 μg/mL lincomycin at 25°C at the specified xylose concentrations. 11 μM isopropyl β -D-1-thiogalactopyranoside (IPTG) was added to induce GFP-MreB and the cells were shifted to 37°C and allowed to grow for 2 hours before imaging. Cells of strain BEG202 (δ*tagO)* with GFP-Mbl under a xylose-inducible promoter were grown overnight at 25°C in LB supplemented with 20 mM magnesium chloride and 0.125 mM xylose, and shifted to 37°C for 2 hours before imaging. Cells were placed on cleaned glass coverslips thickness No. 1.5, as described in the next section. 3-6% agar pads were prepared in LB supplemented with 20 mM magnesium chloride, 11 μM IPTG and the desired concentration of xylose. Images were collected for 3 min at 1 or 2 s intervals, as specified.

### Imaging – slide preparation

Coverslips were sonicated in 1 M KOH for 15 min, followed by 5 washes with water. Coverslips were washed twice with 100% ethanol, and then sonicated in 100% ethanol, followed by one more wash in 100% ethanol. They were stored in ethanol and dried for 10 min before use.

### Imaging – spinning disk confocal

Images were collected on a Nikon TI microscope with a Hamamatsu ImagEM (EM-CCD) camera (effective pixel size 160 nm) and Nikon 100X NA 1.45 TIRF objective. Z stacks were obtained at 0.2 μm slices. Total image depth was 3 μm. Only the top 3 slices of the cell were used in maximum intensity projections in Figure 3D.

### Image processing

All image processing unless otherwise specified was performed in FIJI (Schindelin et al., 2012; Schneider et al., 2012). Images used for particle tracking were unaltered, except for trimming five pixels from the edges of some videos to remove edge artifacts detected by the tracking software. Phase contrast images and fluorescent images of protoplasts were adjusted for contrast. Phase contrast images presented in the manuscript collected from cells in the custom microfluidic device, which did not undergo quantitative processing, were gamma-adjusted (γ=1.5) to compensate for changes in brightness occurring at the device’s feature borders; such processing was not used for growth quantification. The images for Supplementary Movie 2 were background-subtracted for viewing purposes; unaltered images were used for quantitative processing in all cases.

### Microfluidics

The custom microfluidic setup used to confine cells in Figure 3A-C, Supplementary Figure 3A, and Supplementary Movie 3 was previously described in (Norman et al., 2013). Briefly, a polydimethylsiloxane slab with surface features was bonded to a 22 x 60 mm glass coverslip by oxygen plasma treatment followed by heating to 65°C for >1 hr. The features in our setup differed from those described in (Norman et al., 2013), particularly in the omission of a second, wider layer in the cell chambers, which enhanced growth at timescales beyond that of our experiments. Syringes containing growth medium were connected to the microfluidic features using Tygon tubing stainless steel dispensing needles (McMaster Carr Supply Company, Elmhurst, Illinois). Medium was supplied to cells at a constant rate of 2-5 μL/min using automatic syringe pumps. Imaging was carried out using phase contrast microscopy as described above. For the microfluidics experiments in Figures 5 and 6 and Supplementary Movie 6 (top), 7, and 8, the CellASIC platform from Merck Millipore (Billerica, Massachusetts) was used with B04A plates.

### Cell confinement experiments

The cell confinement experiment in Figure 3A-C was conducted by first loading cells into the chamber: BEG300 cells were grown to stationary phase (OD_600_ 3.0 – 5.0) in LB supplemented with 20 mM magnesium chloride, passed through a 5 μm filter, and concentrated 100-fold before loading in the custom-made microfluidic device. Both phase contrast and fluorescent imaging were performed as described in the “Imaging” section above. For observing MreB movement, MreB-GFP expression was induced with 50 μM IPTG upon loading into the microfluidic chamber, and cells were imaged every 2 s with a camera exposure time of 300 ms.

### MreB alignment within protoplasts

Cells of strains bJS18 (GFP-Mbl) and bEG300 (GFP-MreB) were grown overnight at 25°C in the osmoprotective SMM media (LB supplemented with 20 mM magnesium chloride, 17 mM maleic acid, 500 mM sucrose, brought to a pH of 7.0) with maximum xylose induction (30 mM); cells were shifted to 37°C in the morning. For strain bEG300, the SMM media was supplemented with 8mM xylose (for intermediate TagO induction). Following 2 hours of growth, 10 mg/mL of freshly suspended lysozyme was added to the cultures with OD_600_ > 0.2. After growing for 1-2 hours in lysozyme, the cells were spun and concentrated. 6% agar pads made in LB-SMM were made using a polydimethylsiloxane (PDMS) mold with crosses (2, 4 and 5 μm arms and 5 μm center). The cells were placed on the agar pad for 2 min, allowing the cells to settle in the crosses. The pad was then placed in a MatTek (Ashland, Massachusetts) dish for imaging. To check for the presence of cell walls in protoplasts, wheat germ agglutinin conjugated to Alexa-555 was used. 20 μL of 1 mg/mL stock was added to 1 mL of cells 20 min before the start of imaging. Some cultures, after inoculation in the MatTek dish, were incubated at 37°C for 30 min to allow cell growth.

### Depletions in liquid culture

TagO depletions in Figure 2A were conducted using strain BEG300 in liquid culture. Cells were prepared as overnights, as described above, then grown at the specified xylose concentration at 37°C in LB with 20 mM magnesium chloride for 4 hrs. The cells were then imaged as described above in the “Imaging – MreB particle tracking” section.

Pbp2A depletions shown in Figure 2C were conducted in liquid culture using strain

BRB785 with an IPTG-inducible Pbp2A fusion at the native locus with the redundant transpeptidase PbpH deleted. This strain was grown overnight in the presence of 2 mM IPTG, and then inoculated into CH media containing 2 mM IPTG, 0.015% xylose, and 20 mM magnesium chloride to stabilize the cells against lysis. At an OD_600_ of 0.6, cells were spun down in a tabletop centrifuge and washed 3 times in CH media lacking IPTG. Cells were placed under agar pads containing 20 mM magnesium chloride, and spinning disk confocal images were taken every 5 s on a Nikon Ti microscope with a 100X 1.49 TIRF objective and a Hamamatsu ImagEM C9100-13 EM-CCD camera (effective pixel size of 160 nm).

### Depletions under solid state medium

Depletions shown in Figure 5A were conducted using strain BEG300. Cells were prepared as overnights in LB with 1 mM magnesium chloride and 12 mM xylose. In the morning, they were washed in LB with 12 mM xylose and no magnesium and placed under a 3% agar pad with the same medium. Phase contrast images were collected every 5 min using a Photometrics (Tucson, Arizona) CoolSNAP HQ2 CCD camera.

### Repletions

Repletions of TagO or Pbp2a on pads, as shown in Figure 5B and Supplementary Movie 6 (bottom), were performed with strains BEG300 and BRB785 respectively. Cells were grown as overnights, as described above, then depleted at 37°C for >4 hours in LB with 20 mM magnesium chloride and collected by centrifugation at 6,000 x *g* for 2 min. The cells were re-suspended in LB supplemented with 20 mM magnesium chloride and 1 mM IPTG (BRB785) and 30 mM xylose (BEG300), placed under 5% agarose pads on coverslips with thickness No. 1.5 for imaging. Phase contrast images were collected every 5 min using a Photometrics CoolSNAP HQ2 CCD camera.

For the repletions shown in Figures 5B-C, 6A, Supplementary Figure 5C, and Supplementary Movie 6 (top) and 7, performed in the CellASIC microfluidic device in a B04A plate, BCW82 and BEG300 cells were grown to OD_600_ 1.2 – 1.5 in LB supplemented with 20 mM magnesium chloride, centrifuged to pellet large clumps for 3 min at < 500 x *g*, and the supernatant loaded into the plate. Growth medium was supplied at 5-6 PSI. Cells were grown for at least an additional 30 min before the addition of inducer to the growth medium. Phase contrast images were collected every 10 min. Fluorescent images were collected on the imaging setup described in the “Imaging – MreB Particle Tracking” section above: GFP-MreB was induced upon loading into the microfluidic chamber with 1 mM IPTG, and MreB dynamics were observed for 3 min after every 10 min, using 300 ms camera exposures taken every 2 s. For the repletions shown in Figure 6C and S5E, the same procedure was used, but with imaging performed on the spinning disk confocal microscope described in “Imaging – Spinning Disk Confocal”. Z-stacks were collected with a range of 3 μm around the focal plane and 0.2 μm steps. The MreB localization experiments were done using strain bEG300 with full induction of GFP-MreB (1mM IPTG) and recovering cells were imaged using the spinning disk microscope, collecting Z-stacks as described before.

Where indicated, instead of visualizing MreB dynamics, fluorescent D-amino acids (Kuru et al., 2012) (7 μM) were added to the growth medium in the CellASIC device: HADA during depletions of TagO (0 mM xylose) and Cy3B-ADA during repletion of TagO (30 mM xylose). Cells were washed with LB supplemented with 20 mM magnesium chloride containing no D-amino acids for 1-2 min before imaging.

To test if rod shape recovery occurs in the absence of cell division, 3 strains were tested (BAB327, BAB343 and BAB388). Cells of BAB327 and BAB388 were grown in CH media with 25 mM magnesium chloride in the absence of xylose at 37°C until OD_600_ ∼0.5 and diluted 10-fold in fresh media. After 2 hours of growth, IPTG was added to a final concentration of 1 mM (MinCD and FtsA, respectively) and cells were incubated for an extra 1 hour. Cells were imaged on a spinning disk confocal under pads with 1 mM IPTG and 60 mM xylose (for TagO repletion). Phase-contrast and fluorescent images were acquired at 10 minute intervals for a total of 8 hours. Cells of BAB343 were grown in LB supplemented with 20 mM, magnesium chloride in the absence of xylose at 25°C overnight. The next day, after 2 hours of growth in the same media at 37°C, IPTG was added to a final concentration of 1 mM (MciZ) and cells were incubated for an extra 1 hour. Cells were imaged on a spinning disk confocal under pads with 1 mM IPTG and 30 mM xylose (for TagO repletion). Phase-contrast and fluorescent images were acquired at 10 minute intervals for a total of 4 hours.

### Depletion and repletion of Magnesium in the CellASIC

For Supplementary Movie 8, cells of BCW51 were grown overnight at 25° C in LB supplemented with 8mM xylose, 20 mM magnesium chloride, 1 μg/ml erythromycin and 25 μg/ml lincomycin (MLS). Cells were shifted to 37°C for 2 hours and loaded into the CellASIC B04A plate at OD_600_ ∼0.6. At the start of imaging, magnesium was depleted by flowing in LB supplemented only with 8 mM xylose and MLS at 3 psi. Images were collected every 20 min over a 4 hr period. Magnesium was resupplied to the cells by changing to LB supplemented with 8 mM xylose, 20 mM magnesium chloride, and MLS. Imaging was continued every 20 min for an additional 4 hrs.

### Measurements of cell shape at steady state growth

Cells were grown overnight at 25°C in LB supplemented with 30 mM xylose, 20 mM magnesium chloride, 1 μg/mL erythromycin and 25 μg/mL lincomycin. In the morning they were collected at OD_600_∼0.2, spun in a tabletop centrifuge at 9000 rpm for 3 min and washed in LB supplemented with various xylose (0-30 mM) and magnesium chloride (0-20 mM) levels. 25-fold serial dilutions into LB supplemented with the same xylose and magnesium chloride concentrations were made and allowed to grow at 37°C for 4 hrs. Cells at OD_600_ ∼0.2 were concentrated by spinning in a tabletop centrifuge at 9000 rpm for 3 min. They were placed on a coverslip thickness No. 1.5 under 3% agarose pads made in LB supplemented with the same concentrations of xylose and magnesium chloride. Images were collected using the imaging setup described in the “Imaging – phase contrast microscopy” section above, as well as with a Photometrics CoolSNAP HQ2 CCD camera. The magnification and pixel size were the same in both setups.

### Muropeptide analysis

To prepare muropeptides from strain BEG300, a similar protocol was performed as reported previously (Atrih et al., 1999; Kühner et al., 2014). An overnight culture of BEG300 (2 mL) was centrifuged at 10,000 rpm for 5 min. The cell pellet was subsequently resuspended in 1 ml 0.25% SDS in 0.1 M Tris/HCl (pH 6.8). The mixture was boiled at 100 °C for 20 minutes. After cooling, the suspension was centrifuged at 16,000 rpm for 10 minutes, and the pellet was washed with 1.5 ml H_2_0 and centrifuged again at 16,000 rpm for 10 minutes. The pellet was then washed two more times with water. The pellet was then resuspended in 1 ml H_2_0 and sonicated for 30 minutes. 500 μl of a solution containing DNase (15 μg/ml) and RNase (60 μg/ml) in 0.1 M Tris/HCl (pH 6.8) was added. After shaking at 37 ?C for 2 hours, the enzymes were inactivated at 100 ?C for 5 minutes. The mixture was pelleted at 16,000 rpm and washed with 1 ml H_2_0 twice. Wall teichoic acid was removed from the pellet by resuspending the pellet in 500 μl of 1 M HCl and incubating the mixture at 37 ?C for 4 hours. The pellet was then centrifuged and washed with water at least four times to neutralize the pH to approximately pH 6. Subsequently, the pellet was resuspended in 100 μl of digestion buffer (12.5 mM NaH_2_PO_4_) with 10 μ l of mutanolysin (5 U/mL in H_2_O, from *Streptomyces globisporus*, purchased from Sigma Aldrich, St. Louis, Missouri). Peptidoglycan was digested overnight, shaking at 37 °C. Subsequently, the sample was centrifuged at 16,000 rpm for 10 minutes and muropeptides were reduced by adding 50 μl of sodium borohydride (10 mg/ml, H_2_0) was added. The mixture was incubated for 30 minutes. To quench the muropeptide reduction, 1.4 μl of 20% phosphoric acid was added to adjust the pH to 4. The mixture was then used for LC/MS analysis. High-performance liquid chromatography (HPLC) was carried out on an Agilent Technologies (Santa Clara, California) 1260 Quanternary LC system using a SymmetryShield RP18 5 μM, 4.6 x 250 mm column (Waters, Part No. 186000112). Solvent A was 0.1% formic acid in water; Solvent B was 0.1% formic acid in acetonitrile. At a flow rate of 0.5 mL/min, Solvent B was increased from 0-20% in 100 minutes, held at 20% for 20 minutes, increased to 80% by 130 minutes, held at 80% for 10 minutes, and subsequently reduced to 0% and held at 0% for 10 minutes. To analyze the muropeptide composition, muropeptides were identified by the masses observed under specific peaks in the total ion chromatograms. Similar muropeptides were observed as previously reported for other *Bacillus subtilis* strains (Kühner et al., 2014). To quantify the amount of muropeptides observed, exact masses were integrated using extracted ion chromatograms (EICs). Subsequently, we calculated the percentage crosslinking as previously reported (Glauner et al., 1988). Technical and biological replicates were performed for each strain background.

### Particle tracking

The MATLAB based software uTrack was used for particle tracking (Jaqaman et al., 2008). We used the comet detection algorithm to detect filaments (difference of Gaussian: 1 pixel low-pass to 4-6 pixels high pass, watershed segmentation parameters: minimum threshold 3-5 standard deviations with a step size of 1 pixel) which, at our MreB induction levels gave better localization of the resultant asymmetric particles over algorithms that search for symmetric Gaussians. Visual inspection of detected particles confirmed that most of the particles and none of the noise were being detected. A minimum Brownian search radius of 0.1-0.2 pixels and a maximum of 1-2 pixels was applied to link particles with at least 5 successive frames. Directed motion propagation was applied, with no joins between gaps allowed. Tracks were visualized using the FIJI plug-in TrackMate (Tinevez et al., 2017). For sphere to rod transitions and cells confined in microfluidic channels, movies were processed by subtracting every 8^th^ frame from each frame to remove stationary spots using the FIJI plugin StackDifference before tracking. The tracking was done as described earlier in this section.

### Fluorescent analysis of TagTUV

Strains containing fluorescent fusions to TagT, TagU, and TagV were grown as described in the “Overnight culture growth” section but in CH medium instead of LB. Cells were grown for 3 hours at 37°C before imaging, then collected by centrifugation at 6,000 x *g* for 2 min and re-suspended in CH. Cells were then placed on a glass coverslip thickness No. 1.5 under an agar pad thickness 1 mm made from CH and 1.5% agarose. Timelapse images were collected with TIRF illumination, using continuous 100 ms 488 nm exposures. Epifluorescent illuminated images were collected from a single exposure, while maximal intensity projections were formed from a series of continuous 100 ms TIRF exposures.

### Teichoic acid labeling with Concanavalin A

BEG300 cells were grown from overnights as described in the “Fluorescent analysis of TagTUV” section at 37°C for 4 hours without xylose to deplete WTAs, then induced with 30 mM xylose for 1.5 hours to re-induce WTA expression. Cells were then moved to 25°C for at least 30 min and incubated with 25 μg/mL Concanavalin A conjugated to Alexa Fluor 647. Cells were collected by centrifugation at 6,000 x *g* for 2 min, washed with CH medium, then re-suspended in fresh CH medium. Cells were then placed on a glass coverslip thickness No. 1.5 under an agar pad thickness 1 mm made from CH medium and 1.5% agarose. For PY79 and BCW61 controls, lectin-Alexa Fluor conjugate concentration was 200 μg/mL. For non-quantitative analysis, imaging was performed with the microscope described in “Imaging – MreB Particle Tracking”; for quantitative analysis, imaging was performed with the microscope described in “Imaging – Spinning Disk Confocal”. Quantification was performed in FIJI; pixel values were corrected for mean fluorescent background.

### Data analysis – selecting directional tracks

The output of uTrack is the position coordinates of tracks over frames. We fit a line through these coordinates using orthogonal least squares regression to minimize the perpendicular distance of the points from the line of best fit. We used principal component analysis for orthogonal regression using custom written MATLAB code. The R^2^ values we obtain range from 0.5 to 1. We calculated mean track positions, angles and displacement using the line of best fit for all tracks. We also calculated the mean square displacement versus time of individual tracks and fit these curves to the quadratic equation *MSD*(*t*) = 4*Dt* + (*Vt*)2, using nonlinear least squares fitting. As later times have fewer points and are noisier, we fit the first 80% of the data for each track. We determined α by fitting a straight line to the log(*MSD*(*t*)) vs. log (*t*) curve. The goodness of fit was evaluated by determining the R^2^ value. We selected tracks for linearity and directional motion, based on the following cutoffs: R^2^ > 0.9, displacement > 0.2 μm, velocity > 1e^-9^ μm/s, and R^2^ of the linear fit of log(MSD(t)) vs. log(t) > 0.6.

### Data analysis – cell segmentation

The MATLAB-based software Morphometrics (Ursell et al., 2017) was used to segment phase contrast images of cells. We used the phase contrast setting for rod-shaped and intermediate states and the peripheral fluorescence setting for spherical states, because in this latter condition, peripheral fluorescence empirically did a better job of fitting cell outlines. The cell contours obtained were visually inspected and any erroneous contours were removed by custom written MATLAB code.

### Data analysis – track angles with respect to the long axis of the cell

Track angles were calculated with respect to the cell midline as defined by the Morphometrics “Calculate Pill Mesh” feature, which identifies the midline based on a unique discretization of the cell shape determined from its Voronoi diagram. The difference between the track angle and midline angle was then calculated. Since the track angles θ*t* and midline angles θ*m* both ranged from −90° to 90°, the range of angle differencesΔθ = θ*t* - θ*m* was −180° to 180°. We changed the range to 0 to 180° by the transformation: Δθ= 180 + Δθ *if* Δθ< 0, and 0 to 90° by the transformation: Δθ= 180 - Δθ *if* Δθ > 90. The standard deviations (SD) reported are measured from the distributions with a range of 0-180° as this SD most accurately depicts deviations from 90°.

### Data analysis – mean dot product of tracks

Custom written MATLAB code was used to calculate the normalized dot product (*DP*) of track pairs along with the distance (*d)* between their mean positions 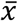 and 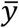 as follows:

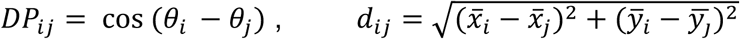

To eliminate out-of-cell tracks we only considered those that had 3 other tracks within a 5 μm radius of their mean position. The dot product of track pairs (DP) and distance (d) between them was stored in data files, along with all the previous information for each individual track (R^2^, velocities, angles, mean positions, displacement etc). The files were then parsed using the cutoffs described in the “Data analysis – selecting directional tracks” section. The tracks were binned based on the distance and the mean dot product calculated for each distance range as follows:

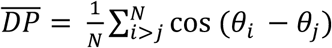

A cutoff of 3 μm was chosen as the maximum binning distance, which is the average length of a cell.

### Data analysis – simulation of random angles

A data file containing simulated tracks was created by a custom written MATLAB script, which generates random angles distributed randomly on a 100 x 100 μm area. Each track has R^2^ = 0.95, velocity = 25 nm/s and displacement = 1 μm. The same analysis code was run on these simulated tracks to generate track pairs with dot product and distance stored in a new data file. The data file was parsed using the same cutoffs as the real data and the mean dot product for each distance range calculated. The total numbers of trajectories within the simulation were much higher than the actual data (2-10 times higher).

### Data analysis – cell width

Pill meshes were created using Morphometrics (Ursell et al., 2017), which calculates the coordinates of line segments perpendicular to the cell long axis. For cell widths at various steady state TagO and Mg^2+^ levels, the distance of these line segments were calculated using a custom written MATLAB script and the maximum width along the length of the cell was taken as the cell width. When measuring cell width nearest to a track (for calculating track angle as a function of cell width), the mean width of the 10 nearest contour points from the track was calculated using a custom written MATLAB script. Cell widths of emerging bulges and rods from round cells were measured manually in FIJI. Our ability to segment individual spherical cells was limited by their nonuniform contrast, perhaps arising from the nonuniform thickness of these cells in the Z dimension; consequently, Morphometrics-based width measurements in these cells was limited, especially in cells exceeding 2 μ m in diameter.

### Data analysis – cell curvature

Sidewall curvature of cells was extracted from the pill mesh obtained from Morphometrics. The curvature values are calculated from 3 successive contour points and smoothed over 2 pixels. The mean curvature of 3 nearest points to each track were calculated from both sides of the cell contour and called the mean curvature. Principal curvature ratio was calculated by dividing the sidewall curvature with the curvature in the radial direction (calculated from cell width assuming the cell is radially symmetric). For radial curvature we used the following expression, where *r*_cell_ is half the cell width:

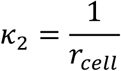

A value close to 1 indicates the two principal curvatures are similar and the cells are round.

### Data analysis – time and curvature plots of rod shape recovery

Phase contrast images were used to show rod shape emergence from local bulges. Edges were enhanced in FIJI and contrast adjusted to give bright cell outlines in the images. The stack was then colored in time using temporal color code function in FIJI. To create the curvature plot, the phase contrast images were run through Morphometrics which calculates the curvature at each contour point along the cell outline. The contour points of interest were selected and plotted using a custom written MATLAB script, which colored each point according to its local curvature as calculated by Morphometrics. To provide a good resolution for positive curvatures, we rescaled the color map such that negative curvatures were colored blue and positive curvatures were scaled by their curvature value.

### Data analysis – single cell doubling times

Data from agar pads experiments was analyzed using custom written MATLAB code. Data from cellASIC experiments was analyzed in Morphometrics to get areas for each cell. For doubling times during sphere to rod transitions, the data was collected by manually measuring the areas of the sphere and rod regions of the same cell in FIJI. In all cases, the area of each cell per frame was calculated and the log plot of area vs time was fit to a line. The doubling time was calculated using the slope of this line.

### Data analysis – Tangential correlation of cell contours

Cell contours were used to calculate tangent angles using the equation: 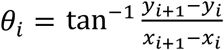 The correlation between angles was calculated using the cosine of the angle difference binned as a function of number of points (n) between the angles: 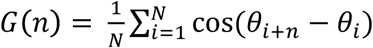The number of points was converted to contour length using the pixel size of the camera to get the final correlation function: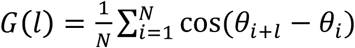

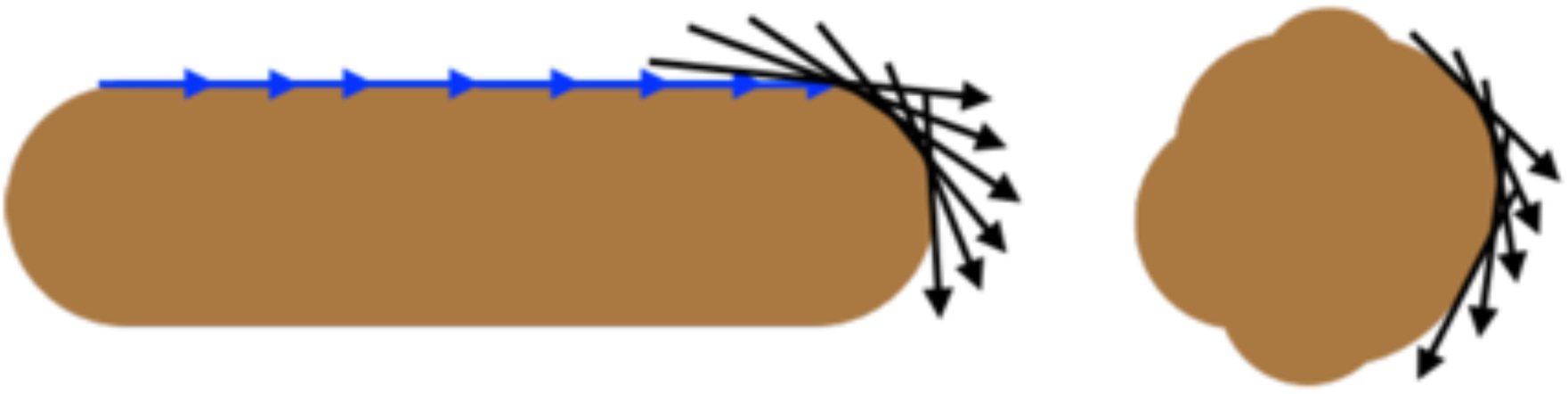

As shown above, for straight rods, the contour angles on average remain highly correlated over larger distances, becoming uncorrelated at the cell pole. In spherical cells, the angles become uncorrelated at shorter distances.

### *T. maritima* MreB protein purification

Full length, un-tagged *Thermotoga maritima* MreB was purified as described previously (Salje et al., 2011).

### *In vitro* reconstitution of *T. maritima* MreB filaments inside liposomes

The protein was encapsulated inside unilamellar liposomes following a previously published protocol (Szwedziak et al., 2014). For this, 50 μL of *E. coli* total lipid extract, dissolved in chloroform at 10 mg/mL, was dried in a glass vial under a stream of nitrogen gas and left overnight under vacuum to remove traces of the solvent. The resulting thin lipid film was hydrated with 50 μL of TEN100 8.0 (50 mM Tris/HCl, 100 mM NaCl, 1 mM EDTA, 1 mM NaN_3_, pH 8.0), supplemented with 20 mM CHAPS (Anatrace, Maumee, Ohio), and shaken vigorously at 800 rpm using a benchtop micro centrifuge tube shaker for 2 hrs. The lipid-detergent solution was then sonicated for 1 min in a water bath sonicator. Subsequently, 50 μL of MreB protein solution at 30 μM, supplemented with 0.5 mM magnesium ATP (Jena Bioscience, Germany) was added and left for 30 min at room temperature. Next, the mixture was gradually diluted within 10-20 min to 600 μL with TEN100 8.0 plus 0.5 mM magnesium ATP (without detergent) to trigger spontaneous liposome formation. 2.5 μL of the solution was mixed with 0.2 μ

L 10 nm IgG immunogold conjugate (TAAB, UK) and plunge-frozen onto Quantifoil R2/2 carbon grid, using a Vitrobot automated freeze plunger (FEI Company, Hillsboro, Oregon) into liquid ethane.

### Electron cryomicroscopy and cryotomography

2D electron cryomicroscopy images were taken on an FEI Polara TEM (FEI Company) operating at 300 kV with a 4k x 4k Falcon II direct electron detector (FEI Company) at a pixel size of 1.8 Å. For electron cryotomography, samples were imaged using an FEI Titan Krios TEM (FEI Company) operating at 300 kV, equipped with a Gatan imaging filter set at zero-loss peak with a slit-width of 20 eV. A 4k x 4k post-GIF K2 Summit direct electron detector (Gatan, a subsidiary of Roper Technologies, Lakewood Ranch, Florida) was used for data acquisition with SerialEM software (Mastronarde, 2005) at a pixel size of 3.8 Å at the specimen level. Specimens were tilted from −60° to +60 ° with uniform 1° increments. The defocus was set to between 8 and 10 μm, and the total dose for each tilt series was around 120-150 e/Å^2^.

### Image processing

Tomographic reconstructions from tilt series were calculated using RAPTOR (Amat et al., 2008) and the IMOD tomography reconstruction package followed by SIRT reconstruction with the TOMO3D package (Agulleiro and Fernandez, 2011; Kremer et al., 1996). Movies showing liposomes were prepared with Chimera and PyMOL (DeLano, 2002; Pettersen et al., 2004).

## Strain construction

**BCW51** [*ycgO::Pxyl-tagO, tagO::erm, amyE::sfGFP-mreB, sinR::phleo*] was generated by transforming BEG300 with a Gibson assembly consisting of three fragments: 1) PCR with primers Sinr_up_F and Sinr_up_R and template PY79 genomic DNA; 2) PCR with primers oJM028 and oJM029 and template plasmid pWX478a (containing *phleo*); 3) PCR with primers Sinr_DOWN_R and Sinr_DOWN_F and template genomic DNA.

**BCW61** [*tagE::erm*] was generated by transforming PY79 with a Gibson assembly consisting of three fragments: 1) PCR with primers oCW054 and oCW055 and template PY79 genomic DNA; 2) PCR with primers oJM028 and oCW057 and template plasmid pWX467a containing *cat*; 3) PCR with primers oCW058 and oCW059 and template PY79 genomic DNA.

**BCW72** [*yvhJωPxylA-mazF (cat)*] was generated by transforming PY79 with a Gibson assembly consisting of three fragments: 1) PCR with primers oCW139 and oCW141 and template PY79 genomic DNA; 2) PCR with primers oJM029 and oMK047 and template DNA consisting of a fusion of *cat* and the *mazF* counterselectable marker from pGDREF (Yu et al., 2010); 3) PCR with primers oCW142 and oCW143 and template PY79 genomic DNA.

**BCW77** [*ywtFωPxylA-mazF (cat)*] was generated by transforming PY79 with a

Gibson assembly consisting of three fragments: 1) PCR with primers oCW159 and oCW161 and template PY79 genomic DNA; 2) PCR with primers oJM029 and oMK047 and template DNA consisting of a fusion of *cat* and the *mazF* counterselectable marker from pGDREF (Yu et al., 2010); 3) PCR with primers oCW164 and oCW165 and template PY79 genomic DNA.

**BCW78** [*ywtFωmsfGFP-ywtF*] was generated by transforming BCW77 with a Gibson assembly consisting of three fragments: 1) PCR with primers oCW160 and oCW161 and template PY79 genomic DNA; 2) PCR with primers oCW072 and oCW073 and BMD61 genomic DNA; 3) PCR with primers oCW163 and oCW165 and template PY79 genomic DNA.

**BCW79** [*yvhJωmsfGFP-yvhJ*] was generated by transforming BCW72 with a Gibson assembly consisting of three fragments: 1) PCR with primers oCW139 and oCW146 and template PY79 genomic DNA; 2) PCR with primers oCW072 and oCW073 and BMD61 genomic DNA; 3) PCR with primers oCW143 and oCW145 and template PY79 genomic DNA.

**BCW80** [*lytRωPxylA-mazF (cat)*] was generated by transforming PY79 with a Gibson assembly consisting of three fragments: 1) PCR with primers oCW101 and oCW109 and template PY79 genomic DNA; 2) PCR with primers oJM029 and oMK047 and template DNA consisting of a fusion of *cat* and the *mazF* counterselectable marker from pGDREF (Yu et al., 2010); 3) PCR with primers oCW100 and oCW125 and template PY79 genomic DNA.

**BCW81** [*lytRωmsfGFP-lytR*] was generated by transforming BCW72 with a Gibson assembly consisting of three fragments: 1) PCR with primers oCW101 and oCW137 and template PY79 genomic DNA; 2) PCR with primers oCW072 and oCW073 and BMD61 genomic DNA; 3) PCR with primers oCW100 and oCW138 and template PY79 genomic DNA.

**BCW82** [*tagO::erm*, *ycgO::PxylA-tagO*, *amyE::Pspac-gfp-mreB (spec)*, *dacA::kan*] was generated by transforming BEG300 with genomic DNA from BGL19.

**BEG202** [*tagO::erm amyE::Pxyl-gfp-mbl (spec)*] was generated by transforming BEB1451 with genomic DNA from BJS18.

**BEG281** [*ycgO::PxylA-tagO*] was generated by transforming with a plasmid created via ligating a Gibson assembly into pKM077. pKM77 was digested with EcoRI and XhoI. The assembly was created with two fragments: 1) PCR with primers oEG85 and oEG86 and template py79 genomic DNA; 2) PCR with primers oEG87 and oEG88.

**BEG291** [*tagO::erm*, *ycgO::PxylA-tagO]* was generated by transforming BEG281 with genomic DNA from BRB4282.

**BEG300** [*tagO::erm*, *ycgO::PxylA-tagO*, *amyE::Pspac-gfp-mreB (spec)*] was generated by transforming BEG291 with genomic DNA from BEG275.

**BMD61** [*mblωmbl-msfGFP (spec)*] was generated by transforming py79 with a Gibson assembly consisting of four fragments: 1) PCR with primers oMD44 and oMD90 and template PY79 genomic DNA; 2) PCR with primers oMD47 and oMD56 and template synthetic, codon-optimized *msfGFP*; 3) PCR with primers oJM028 and oJM029 and template plasmid pWX466a (containing *spec*); 4) PCR with primers oMD48 and oMD50 and template genomic DNA.

**bSW99** [*amyE::spc-Pspac-mciZ*] was generated by transforming PY79 with a Gibson assembly consisting of five fragments: 1) PCR with primers oMD191 and oMD108 and template PY79 genomic DNA (containing upstream region of amyE); 2) PCR with primers oJM29 and oJM28 and template plasmid pWX466a (containing *spec*); 3) PCR with primers oMD234 and oSW76 and template plasmid pBOSE1400 (a gift from Dr. Briana Burton, containing *spec*); 4) PCR with primers oAB307 and oAB291 and template PY79 genomic DNA (containing mciZ); 5) PCR with primers oMD196 and oMD197 and template PY79 genomic DNA (containing downstream region of amyE).

**bAB343** [*tagO::erm, ycgO::cat-PxylA-tagO, amyE::spc-Pspac-mciZ, ftsAZ::ftsA-mNeonGreen-ftsZ*] was generated by transforming bAB185 (Bisson-Filho et. al, 2017) with genomic DNA from bSW99*. The resultant strain was then transformed with the genomic DNA from BEG291 and selected for Cm resistance. Subsequently, the resultant strain was transformed again with genomic DNA from BEG291, but colonies were selected for MLS resistance in the presence of 30 mM of xylose and 25 mM* MgCl2.

**bAB327** [*tagO::erm, ycgO::cat-PxylA-tagO, amyE::spc-Physpank-minCD, ftsAZ::ftsA-mNeonGreen-ftsZ*] was generated by transforming bAB185 (Bisson-Filho et al., 2017) with genomic DNA from JB60 (a gift from Dr. Frederico Gueiros-Filho)*. The resultant strain was then transformed with the genomic DNA from BEG291 and selected for Cm resistance. Subsequently, the resultant strain was transformed again with genomic DNA from BEG291, but colonies were selected for MLS resistance in the presence of 60 mM xylose and 25 mM* MgCl2.

**bAB388** [*tagO::erm, ycgO::cat-PxylA-tagO, amyE::spc-Physpank-ftsA, ftsAZ::ftsA-mNeonGreen-ftsZ*] was generated by transforming bAB199 (Bisson-Filho et al., 2017) with genomic DNA from *BEG291 and selected for Cm resistance. Subsequently, the resultant strain was transformed again with genomic DNA from BEG291, but colonies were selected for MLS resistance in the presence of 60 mM xylose and 25 mM* MgCl2.

## Supplemental Text 1 - MreB Modeling

### Modeling predicts preferred MreB orientation and a typical cell width for losing binding orientation

Here we show that energetic modeling of an MreB filament directly binding to the inner membrane predicts the existence of both a preferred orientation of binding and a typical cell width for losing binding orientation. MreB monomers assemble into higher-order oligomers and bind directly to the inner membrane. When an MreB filament binds to the inner membrane, the combined MreB-membrane system requires an energy of deformation *E*_*def*_(*l*_*b*_) for the membrane to deviate from an equilibrium position and gains an energy of interaction *E*_*int*_(*l*_*b*_) from the hydrophobic binding. Both the deformation and interaction energies are expressed as functions of the bound MreB length, *l*_*b*_. Note that the rigid cell wall imposes a boundary constraint on the cell membrane and that the equilibrium membrane configuration arises from a balance of membrane bending, turgor pressure, and cell wall confinement. If the MreB filament were to bind, the change in the total energy *E* of the membrane-MreB system is:

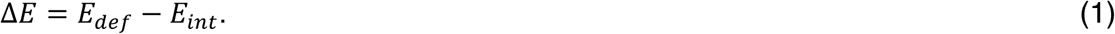

The binding configuration that minimizes Δ*E* corresponds to the one that is observed physically. We therefore wish to minimize Δ*E*.

### Estimate of the hydrophobic interaction energy *E*_*int*_

We assume that the biochemistry of MreB is conserved in prokaryotes so that, like *C. crescentus* and *E. coli* MreB (van den Ent et al., 2014), *B. subtilis* MreB is assembled into antiparallel double protofilaments consisting of many monomeric units. Consider an MreB filament containing *N*_*int*_ interaction sites with a membrane, each with some independent and additive interaction energy 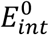. Due to the antiparallel arrangement of the protofilaments (Salje et al., 2011; van den Ent et al., 2014), there are two binding sites per monomeric unit of MreB. We therefore estimate the number of binding sites per MreB binding length *l*_*b*_as:

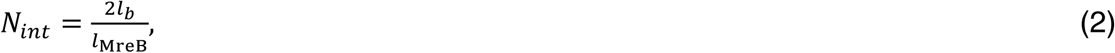

where *l*_MreB_ ∼ 5.1 nm is the length of a monomeric unit. The energy of burying the amino acids relevant to the binding is approximately:

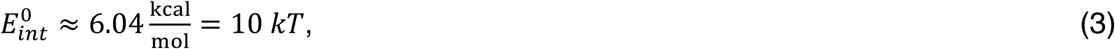

where *k* denotes Boltzmann’s constant and *T* denotes the ambient temperature; the energies of burying individual amino acids were derived from water/octanol partitioning. At a room temperature of *T* = 25*°*C, the interaction energy per MreB binding length *l*_*b*_ is therefore:

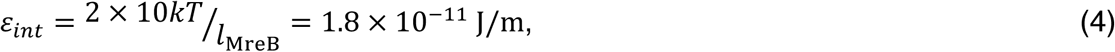

and the hydrophobic interaction energy is: 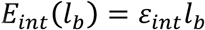

### Estimate of the membrane deformation energy

The membrane deformation energy *E*_*def*_(*l*_*b*_) can be decomposed as:

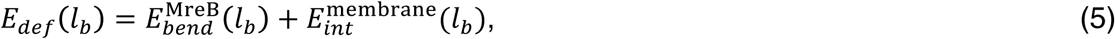

where 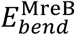 denotes the bending energy of the MreB filament and 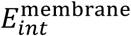 denotes the indentation energy of the membrane. We wish to find the MreB-membrane configuration that minimizes the sum of these terms. We prescribe the forms of these terms as follows.

### The bending energy of an MreB filament

We model an MreB filament as a cylindrical rod, with circular cross-sections of radius *r*_MreB_ and an intrinsic curvature 1/*R*_MreB_ The elastic energy density per unit length of bending a cylindrical rod of cross-sectional radius *r*_MreB_ from a curvature of 1/*R*_MreB_ to a curvature of 1/*R* is given by:

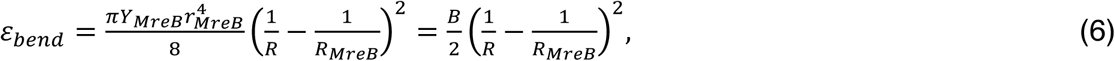

where *Y* _MreB_ is the Young’s modulus of an MreB filament and *B* is its flexural rigidity (Landau and Lifshitz, 1970). Assuming the Young’s modulus of actin, we note that *B* = 1.65 × 10^-25^j ·m which is two orders of magnitude smaller than that previously assumed by Wang and Wingreen for an MreB bundle of cross-sectional radius 10 nm (Wang and Wingreen, 2013). In particular, we assume that MreB binds to the inner membrane as pairs of protofilaments and does not bundle. For a uniform flattening of the MreB filament corresponding to *R* = ∞, *ε*_*bend*_ ≤ 1.65 × 10^-25^ which is less than the MreB-membrane interaction energy *ε*_*int*_ computed above. This suggests that an MreB filament may be susceptible to bending at our energy scale of interest. How much the MreB bends is determined by a trade-off between the polymer bending energy and the indentation energy of the membrane, which we discuss next.

### The membrane Hamiltonian

We model the inner membrane as an isotropic, fluid membrane composed of a phospholipid bilayer, where there is no in-plane shear modulus and the only in-plane deformations are compressions and expansions. The membrane indentation energy can be expressed as the minimum of an energy functional over the indented states of the membrane. This functional is given by the Helfrich Hamiltonian:

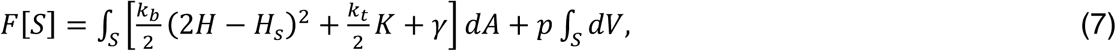

where *k*_*b*_ is the bending rigidity of the membrane, *k*_*t*_ is the saddle-splay modulus of the membrane, *H*_*s*_ is the spontaneous curvature of the bilayer, *γ* is the membrane surface tension, *p* is the pressure differential at the membrane interface, and *H* and *K* are the mean and Gaussian curvatures of the surface *s*, respectively (Safran, 2003; Zhong-Can and Helfrich, 1989). The bending rigidity *k*_*b*_, which depends on membrane composition, is typically 10 to 20 kT for lipid bilayers (Phillips et al., 2012). Assuming that phospholipids are in excess in the bulk and rearrange themselves on the membrane surface to accommodate areal changes (Safran, 2003), we take the membrane surface tension *γ* = 0. For large deformations of the inner membrane such as those induced by cell wall lysis (Deng et al., 2011), the assumption that the phospholipids are in excess in the bulk may fail to hold and result in a nonzero surface tension. A nonzero surface tension would only enhance the energetic preference of the correct binding orientation; hence, taking a finite surface tension would not change our conclusions. The mechanical energy needed to deform the membrane is the difference between the free energies in the deformed *S* and undeformed *S*_0_states:

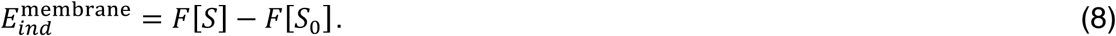

The surface integrals of the Gaussian curvature are topological invariants by the Gauss-Bonnet theorem and therefore cancel in the difference, hence:

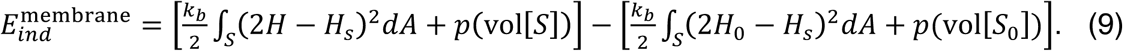

Here *H* denotes the mean curvature of the state *S* and *H*_*0*_ denotes the mean curvature of the state *S*_*0*_. For simplicity, we set the spontaneous curvature *H*_*S*_ = 0; the case of nonvanishing spontaneous curvature can be considered in a similar manner. We therefore write:

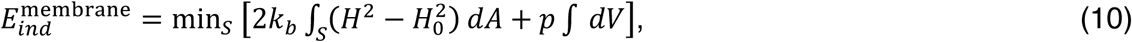

where the volume integral is understood to be the difference of the volumes in the deformed and undeformed states and the areal change accompanying the membrane deformation is small, i.e. *dA*_0_ ≈ *dA* We define the membrane bending energy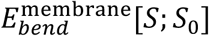 for a conformation *S* to be the former term and the membrane *pV*energy 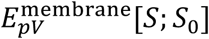to be the latter term in the right hand side of Equation 10.

As *k*_*b*_ is typically 10 to 20 kT, we take *k*_*b*_ = 10 kT and *p* as a parameter of the model. Note that *p* denotes the pressure difference acting on the membrane. The value of *p* is important for determining the tradeoff between the membrane indentation energy 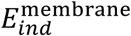 and the MreB bending energy 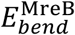accurately, but we will show that the preferred orientation of MreB binding is robust over a broad range of *p*.

### Mechanical equilibrium of the undeformed membrane

Consider the balance of forces on the inner membrane in the undeformed state. Assuming that the undeformed membrane is a cylinder with radius *r* and length *L* and that sufficient phospholipids exist in the bulk so that *γ* = 0, the membrane free energy is

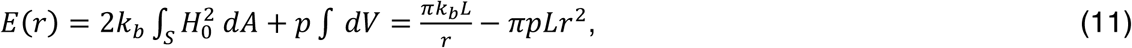

which is monotonically decreasing in *r*This implies that the membrane radius should be maximal at equilibrium. If *p* = 0, then the membrane should press against the cell wall and squeeze out the periplasmic space due to minimization of the bending energy. A model in which the periplasm and cytoplasm are isosmotic (Sochacki et al., 2011) with no mechanical force exerted by the periplasm is therefore inconsistent with the existence of a periplasm. For the periplasm to exist at equilibrium, it must contribute an additional energy term *E*_*peri*_ to the total energy, so that the total energy *F* = *E* + *E*_*peri*_ as a function of *r* has a stable fixed point at *r*_0_ = *r*\*+ *δr*\* Here we define *r*\* = *R*_*cell*_ × *h*_*peri*_, where *R*_*cell*_ is the radius of the cell and *h*_*peri*_ is the thickness of the periplasm, and *δr∗* as the initial deformed height where *F*’(*r*) = 0|_*r*=*r*_0__ We consider expansions of *E*_*peri*_ and *F*(*r*) around *r*_0_ As a function of the deviation in membrane height *δr* = *r* ×*r*_0_ we take *E*_*peri*_ ≈ *κL*(*δr*)^2^ and

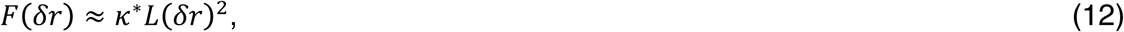

where *k∗* = *k* - *pπ* is the effective membrane pinning modulus, which has been examined before in Wang and Wingreen’s work (Wang and Wingreen, 2013). For the stability of the fixed point at *r*_*0*_, the condition that the second derivative *F″*(*δr*) is positive at *δ* = 0 implies that. However, the validity of the expansion *E*_*peri*_(*δr*) = κ*L*(*δr*)^2^ may be questionable when the deformed height due to polymer binding is larger than or comparable to *δr*\* ∼ *pr*/κ. Hence, the pinning model may be invalid when *p* is vanishingly small. For various combinations of *k* and the polymer bending rigidity where this double-bind is avoided, such as that assumed by Wang and Wingreen’s model, the periplasm is effectively a rigid body. In this case, although a pinning potential can self-consistently penalize deviations in membrane height, it is more intuitive to take the formal limit κ ⟶ ∞ and treat the periplasm as undeformable. We therefore model the periplasm as a rigid, undeformable body that mechanically supports the cell membrane and imposes a boundary condition on the membrane shape. Any deviation from the equilibrium membrane shape induced by MreB binding is then resisted by the full effect of turgor. For this reason, in the following analysis we take *p* = *p*_*cell*_, stipulate that the MreB cannot indent the inner membrane outwards, and do not consider the energetic contribution of *E*_*peri*_

### Configuration with a uniformly bent MreB filament: first-order approximation

With the membrane Hamiltonian as defined in the section above, we now see that the total membrane deformation energy is given by the sum of the MreB bending energy and the membrane indentation energy:

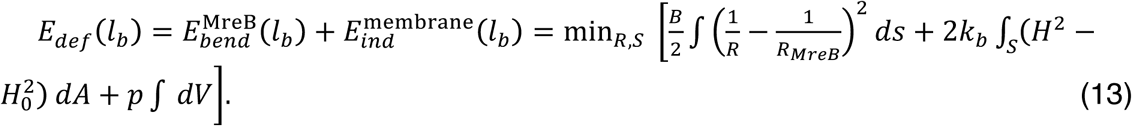

The minimization of equation (13) over all surfaces *s* and MreB curvatures *R*=*R*(*s*) is generally difficult since minimization of the MreB bending energy determines the preferred conformation of MreB, which in turn restricts the set of surfaces *S* that equation (13) must be minimized over. In their work, Wang and Wingreen undertook an elegant approach to minimizing a similar combination of energies by writing the membrane indentation energy in Fourier space. Unlike a membrane pinning term, the pressure-volume energy in equation (13) does not admit a simple Fourier space representation. Nevertheless, considerable insight can be obtained by assuming that MreB bends uniformly. In this case, MreB deforms from a bent cylinder with a native Curvature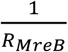 to a bent cylinder with a constant, membrane-bound curvature 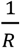In thefollowing, we will take the radius *RR* of the bend to be a parameter in estimating corresponding membrane indentation energy 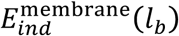will be determined later. We will also assume that MreB binds perpendicular to the cell’s long axis, so that the curvature of the membrane in its undeformed state is simply 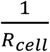, and determine the corrections due to a deviatory binding angle later.

To estimate 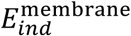, we first examine the energetic contribution of the region *C* of *M* directly involved in the MreB-membrane interaction. Note that the biochemical conformation of MreB, particularly, the antiparallel orientation of its protofilaments, constrains the geometry of the MreB-membrane binding interface. Since we describe the interface *C* as the surface of a bent cylinder with principal radii of curvature 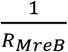 and 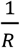, we have:

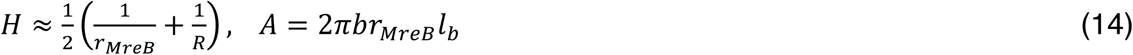

where *r*_*MreB*_ is the cross-sectional radius of MreB and *ππ* is the fraction of interaction along a cross-section of the MreB filament. Thus:

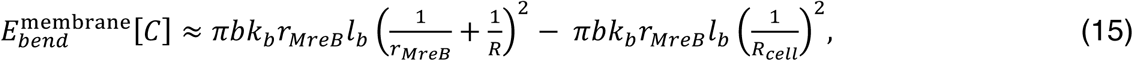

where *H*_k_ = 1/2*R*_*CELL*_ and *R*_*CELL*_ denoting the cell radius, is the mean curvature of the undeformed surface. The contribution of the *pV* energy over *C* can be similarly approximated by finding the area between two circles, one being the MreB filament and the other being the cross-section of a cell with radius *R*_*CELL*_, with *R*_*CELL*_ ≥ *R* as follows:

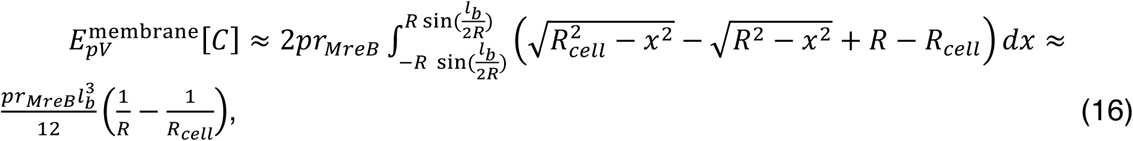

where we have assumed that *l*_*R*_ and 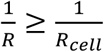 This means that the approximation above is only valid for cases where the MreB filament can only bend up to a curvature 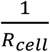. Now, since 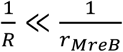, we deduce that the principal bending energy contributionover *C* arises from having the inner membrane tightly wrapped around an MreB filament. For *k*_b_ = 10*kT* and *b* = 1/6, 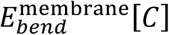 [*C*] takes on a value of:

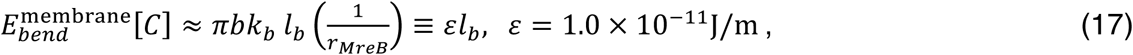

which is smaller than, but comparable in scale to, the interaction energy *E*^int^ computed above. Writing out only the energetics of the binding region *C* under the uniform bending assumption, we therefore see that:

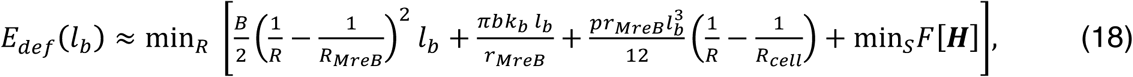

where the last term is the energetic contribution of the falloff region ***H***=*S* − *C* In the case that 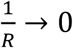, estimates of the values of the first three terms in equation (18), for the parameter values summarized in Table S1, are 10^-19^J, 10^-18^J, and 10^-17^J respectively. This means that, as MreB binds to the inner membrane, the resulting deformation will tend to minimize volumetric changes at the cost of inducing membrane curvature and filament bending. The energetic contribution of the falloff region ***H*** can only be quantitatively accounted for by explicitly finding the membrane shape, which encompasses a tradeoff between the membrane bending energy and the *pV* energy: the former term favors a gradual decay of the indentation, while the latter term prefers a steep decay as to minimize volume. Below, we find that it suffices to consider the case where MreB bends to match the cell curvature: *RR* = *RR*_*CELL*_. In this case, the energetic contribution of the falloff region ***H*** is vanishingly small compared to that of the binding region, since the membrane can heal in a manner in which its mean curvature is small compared to the mean curvature of the binding region. The energetic contribution of ***H*** can therefore be neglected, and we quantify it in future work.

Δ***E* for the pure bending of an MreB filament**

By examining the form of the energetics just over the region *C*, we note that the inclusion of a large pressure *p* increases the energetic preference of an MreB filament binding perpendicular to the cellular long axis.

Consider a case where *p* ≥ *p∗* for some *p∗* to be determined, so that it is energetically unfavorable to displace the membrane volume as opposed to bending the MreB filament. In this case, as discussed above, the energetic contribution of the falloff region ***H*** can be neglected, and an estimate for the minimal value of such a pressure can be obtained by requiring that:

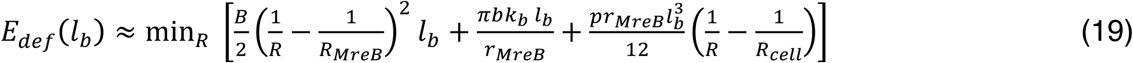

as a function of *RR*, be minimal at *RR*_*CELL*_. For the numerical values relevant to MreB above and summarized in Table S1, this indicates that:

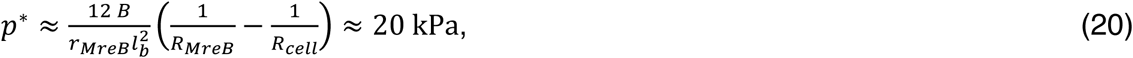

which is 1/100^th^ of the turgor pressure of *B. subtilis*. In this case, assuming *R*=*R*_*cell*_,

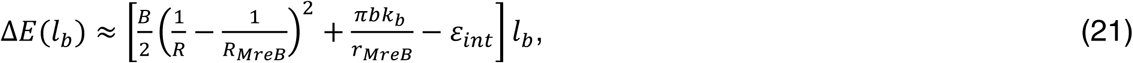

The energetic dependence on *RR*_*CELL*_ is then manifested through the pure bending of MreB when binding to the inner membrane: in particular, we may assume that the MreB filament will always bend to attain a curvature matching that of the cell’s (although small deviations in the membrane height may lead to an even lower energy conformation), and the energetic contribution of the falloff region ***H*** can be neglected.

If *p* < *p∗*, note that both the membrane and the MreB filament can deform each other in a manner that minimizes the total energy, with the membrane shape determined by the geometry of the falloff region ***HH***. For vesicles with a pressure gradient *p* ≈ 0, the fact that an MreB filament grossly deforms the membrane and generates membrane curvature (Salje et al., 2011) is predicted by the shape of ***HH***. For *p* ≈ 0, it can also be shown that the energetic difference between MreB binding at *RR*_*CELL*_ = 500 nm and *RR*_*CELL*_ = 3000 nm is on the order of several *kkkk*, and so a perpendicular alignment of MreB filaments may also be energetically favorable. For simplicity, in the following discussion we shall consider the wild-type cell scenario, where *p* = *p*_*CELL*_ > *p∗*, so that only the pure bending of MreB and the associated membrane bending energy need to be considered in Δ*E*. We will therefore assume the form of equation (21) for Δ*E* in the discussion that follows.

### Preferred orientation of MreB binding

Equations (19) and (21) describe the change in free energy due to MreB binding at perpendicular angle and bending completely to match the curvature 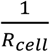 of the cell membrane at this angle. Our modeling then predicts that MreB filaments tend to bind at an angle of θθ = 90*°* relative to the long axis of *B. subtilis*: at any deviatory angle |θθ - 90*°*| > 0*°*, the leading-order correction to the cell radius *R*_*CELL*_ in the MreB bending energy is a multiplicative factor of 1/ cos θθ, which monotonically increases the MreB bending energy. Since the MreB bending energy is minimal when the principal axis of curvature of the MreB filament matches that of the cell and the curvature of the cell is maximal at a perpendicular binding angle, the angle distribution is symmetric about a minimum centered at θθ = 90*°*. This reasoning suggests the existence of a potential well centered at θθ = 90*°*, as shown in Figure 4F under the simplifying assumption that *b* = 0. The depth of this potential well is on the order of tens of *kT*, which appears to be a large enough energetic preference as to be robust to sources of stochasticity such as thermal fluctuations. Furthermore, our discussion shows that membrane binding energetics, and in particular the pure bending of an MreB filament, may complement the conjecture in Salje *et al.*’s work that the membrane insertion loop and amphipathic helix help ensure orthogonal membrane binding (Salje et al., 2011).

A sensitivity analysis shows that the energetic difference of an MreB filament perpendicularly binding to a region of ambient curvature *RR*_*CELL*_ = 0.5 *μ*m and 1.5 *μ*m is still on the order of tens of *kT* over a wide range of binding energy and flexural rigidity values, as shown in Figure 4G. Thus, we anticipate the alignment to be robust to changes in these two parameters.

### A typical cell radius for losing orientation

Our modeling shows that the depth of the potential well in θθ is inversely related to *RR*_*CELL*_, so that at larger cell diameters the angle distribution becomes more uniform. Varying the cell radius *RR*_*CELL*_ from 0.5 microns to 3.0 microns results in a reduction of the depth of the potential well, as illustrated in Figure 4F. We therefore anticipate MreB filaments to bind with more variance in angle for higher values of *RR*_*CELL*_, consistent with the existence of a typical radius at which the binding angle becomes less robust and affected by factors such as thermal fluctuations or other sources of stochasticity.

### Binding orientation at regions of different Gaussian curvatures

Our modeling predicts that, in live cells, MreB filaments will bend to conform to the shape of the inner membrane. Since the binding sites are located at the outer edge of a curved MreB filament, our modeling also predicts that the binding angle distribution becomes more narrow at regions of negative Gaussian curvature: to bind in a conformation that deviates significantly from the preferred binding orientation, in which the filament’s deformed curvature remains of the same sign as its intrinsic curvature, an MreB filament must bend to the extent that its curvature flips sign. Similarly, at regions of positive Gaussian curvature, the binding angle distribution will be less narrow.

Representative binding angle distributions are shown in three cases of positive, vanishing, and negative Gaussian curvatures in Fig. 7B of the main text.

## Supplemental Text 2 – Localization of WTA ligases and WTAs

In order to understand how wall teichoic acid (WTA) depletion and recovery affects cell shape, we observed the spatial localization of WTAs and the extracellular ligases (the genes *lytR*, *yvhJ*, and *ywtF* (Kawai et al., 2011)) that determine their attachment to the cell wall. Previous work has suggested that YwtF and LytR localize in MreB-like patterns (Kawai et al., 2011) and associate with MreB (assayed by *in vivo* crosslinking and tandem affinity purification). We reasoned that if the synthesis or insertion of WTAs is MreB associated, then the emergence of discrete rod-shaped cells upon *tagO* repletion might correlate with preferential WTA insertion at the emerging rod, where MreB shows oriented motion. To test this, we examined 1) the localization and dynamics of fluorescent tagged WTA ligases, and 2) the spatial localization of WTAs in cells recovering from spheres into rods. We constructed sfGFP fusions to each of the WTA ligases under their native promoters, and examined their localization using TIRF and epifluorescent illumination. Although we did observe variation in ligase intensity around the cell periphery under epifluorescent illumination, we did not see any characteristic banding across the cell surface as is seen with MreB using TIRF microscopy (Fig. S6C). Furthermore, TIRF imaging of these fusions at different frame rates did not show any directional motions, indicating they are not moving along with MreB filaments; rather these motions suggested the WTA ligases were diffusing on the cell membrane (Movie S9). We cannot rule out the possibility that the WTA ligases interact with MreB through transient associations.

We next explored the localization of WTAs, using fluorescently labeled Concanavalin A (ConA). ConA is a sugar-binding protein with specific affinity for α-D-glucose, which decorates WTA polymers. ConA has previously been used to localize WTAs (Birdsell et al., 1975; Doyle et al., 1975). The gene *tagE* encodes the glycosylase that adds α-D-glucose to WTA molecules (Allison et al., 2011), which is recognized by ConA. ConA staining of cells deleted for *tagE* shows no staining (Fig. S6A), verifying the specificity of ConA for WTAs over other surface sugars. We then used this probe to examine WTA localizations during recoveries. Addition of ConA to WTA-depleted cells in bulk culture shows very little staining at the cell periphery, consistent with a basal level of WTA expression; induction of *tagO* results in a dramatically increased intensity of staining. Even at early time points, when rod-shaped cells are just starting to appear in the population, WTA staining is relatively uniform, with no patterns reminiscent of the patchy distribution of MreB (Fig. S6B). Together, this data indicates that WTA ligases and WTA incorporation occur uniformly around the cell in both wild type cells as well as in TagO depleted spheres recovering into rods. These findings suggest that the changes in activity of TagO that cause the loss of rod shape (or the reformation thereof) occur uniformly around the cell wall. Furthermore, these findings also suggest that the WTA ligases do not consistently localize to MreB within *B. subtilis*.

## Supplemental Movie Legends

All supplemental movies are available at http://garnerlab.fas.harvard.edu/mreb2017/

**S1. Related to Figure 1C -** Movie showing the trajectories taken by Mbl filaments frequently cross each other close in time. BDR2061, containing GFP-Mbl expressed at the native locus under a xylose-inducible promoter, was induced with 10 mM xylose and imaged with TIRFM. Frames are 1 s apart. Scale bar is 5 μm.

**S2A. Related to Figure 2A –** (***first sequence)*** Timelapse showing circumferential motions of GFP-MreB in rod shaped cells with high TagO expression (BEG300 with 30 mM xylose, and GFP-MreB induced with 50 μM IPTG) ***(second sequence)*** Timelapse of GFP-MreB trajectories in equivalent conditions. ***(third sequence)*** Timelapse showing isotropic motions of GFP-Mbl in a *tagO* knock out strain (BEG202, GFP-Mbl was induced with 0.125 mM xylose). ***(fourth sequence)*** Timelapse of GFP-Mbl trajectories in equivalent conditions as above. Frames are 1 s apart in the first and second sequences, 2 s apart in the third and fourth. All Scale bars are 1 μm.

**S2B. Related to Figure 2C – *(top)*** Timelapse of GFP-Mbl trajectories occurring 2 hours after the initiation of Pbp2a depletion ***(middle and bottom).*** Timelapse of GFP-Mbl trajectories occurring 3 hours after initiation of Pbp2a depletion, where cells become a mixture of rod shaped and round cells. GFP-Mbl shows a mixture of circumferential (bottom) and isotropic (middle) motion. BRB785 was grown in 1 mM IPTG, washed, then grown in media lacking IPTG. Cells were placed under a pad at the indicated times, and imaged with spinning disk confocal. Frames are 5 s apart. Scale bar is 2.5 μm.

**S3. Related to Figure 3A-C -** Timelapse showing circumferential motion of GFP-MreB in BEG300 induced at low TagO levels (2 mM xylose) when confined into long 1.5 x 1.5 μm channels. GFP-MreB was induced with 50 μM IPTG. Frames are 2 s apart. Scale bar is 5 μm.

**S4. Related to Figure 3D-F –** Timelapse of GFP-Mbl in protoplasted cells showing Mbl does not move directionally. BJS18 (containing GFP-Mbl expressed at an ectopic site under xylose control) was induced with 30 mM xylose. Cells were then protoplasted in SMM and grown in molds as detailed in methods. Frames are 1 s apart. Scale bar is 5 μm. Movie was gamma-adjusted, γ = 0.8.

**S5. Related to Figure 4 – *(first sequence***) PyMOL volume rendering of an electron cryotomography 3D map *of T. maritima* MreB included in a liposome (corresponds to liposome depicted in Fig. 4E. (***second sequence)*** Typical field view of an MreB liposome reconstitution experiment. The movie scans through consecutive Z-layers of the tomographic 3D reconstruction. Note that the smaller, round liposomes trapped inside the rod-shaped liposomes are not decorated with MreB filaments.

**S6. Related to Figure 5 – *(top and middle)*** Timelapses showing the local recovery of rod shape upon TagO reinduction from depleted cells. Note the relatively fast growth of rods compared to parent spheres. BEG300 was grown in media lacking xylose, then either loaded into a cellASIC device (top row) or placed under an agar pad (middle row). Both rows were shifted to 30 mM xylose to induce rod-shape recovery, prior to image acquisition. Frames are 10 min apart. Scale bar is 5 μm.

***(bottom)*** Timelapse showing the local recovery of rod shape upon Pbp2a reinduction from cells depleted of Pbp2a/PbpH. BRB785 was grown media lacking IPTG for 4.5 hours, then placed on a pad with 1 mM IPTG before the start of imaging. Frames are 5 min apart. Scale bar is 5 μm.

**S7. Related to Figure 5 and 6 –** Timelapse of rod shape recoveries showing that circumferential MreB-GFP motion A) occurs immediately upon the formation of rod shape, and B) that circumferential motion only occurs in rod-shaped cells, even while attached non-rod cells show unaligned motion. BEG300 was grown overnight in 0mM xylose to deplete TagO. Cells were then loaded into a cellASIC chamber and grown in the same media with 1 mM IPTG to induce GFP-MreB. Prior to imaging, *tagO* expression was reinduced by switching media to contain 30mM xylose. GFP-MreB was imaged with TIRFM. Frames are 2 s apart in the fluorescent channel (green) and 10 min apart in the phase contrast channel (grayscale). Scale bar is 5 μm.

**S8. Related to Figure 5 –** Timelapse showing the loss and recovery of rod shape in cells with intermediate TagO levels when magnesium is removed and added back to the medium. BCW51 was grown in LB supplemented with 8 mM xylose and 20 mM magnesium, then loaded into a cellASIC chamber, and grown in the same media for 30 minutes. At the start of the video the media is switched to contain 0 mM magnesium, causing the cells to lose rod shape. At 4:00:00 the media is switched to contain 20 mM magnesium where the cells revert back into rod-shaped cells. Frames are 20 min apart. Scale bar is 1 μm.

**S9. Related to Figure 6B –** Timelapse showing that the teichoic acid ligases TagTUV do not move circumferentially. Strains shown are BMD61, BCW81, BCW79 and BCW78, where Mbl, TagU (LytR), TagV (YvhJ), and TagT (YwtF) respectively are fused to msfGFP, and expressed from their native promoters. Cells were grown in CH medium and imaged using TIRF illumination every 100 ms. Scale bar is 5 μm.

## Supplemental Tables

**Table S1:** Model Parameters. Related to Figure 4 and Supplemental Text 1.

| Quantity | Estimate | Source |
| --- | --- | --- |
| **MreB values** |  |  |
| MreB bound length $l_b$ | 220 nm | This work |
| MreB monomer length $l_{MreB}$ | 51 angstroms | (van den Ent et al., 2014) |
| MreB cross-sectional radius $r_{MreB}$ | 3.2 nm | (van den Ent et al., 2014) |
| MreB wild-type principal radius of curvature $R_{MreB}$ | 300 nm | This work |
| MreB Young's modulus $Y_{MreB}$ | Similar to actin; 2 GPa | (Kojima et al., 1994) |
| MreB cross-sectional binding fraction $b$ | 0 | This work |
| **Cell values** |  |  |
| *B. subtilis* periplasm thickness $h_{peri}$ | 22 nm | (Matias and Beveridge, 2005) |
| *B. subtilis* cross-sectional radius $R_{cell}$ | 500 nm | This work |
| *B. subtilis* internal turgor pressure $p_{cell}$ | 20 atm | (Whatmore and Reed, 1990) |
| **Binding energy values** |  |  |
| Unit MreB-cell membrane interaction energy $E_{int}^0$ | 10 kT | This work |
| Absolute temperature $T$ | 300 K | This work |
| *B. subtilis* typical cross-sectional radius $R_{cell}^*$ for losing shape | about 1-1.5 microns | This work |

**Table S2:** Strains used in this study.

| Strain | Genotype (all strains are Py79 unless otherwise noted) | Source |
| --- | --- | --- |
| BCW51 | *ycgO::Pxyl-tagO, tagO::erm, amyE::sfGFP-mreB, sinR::phleo* | This work |
| BCW61 | *tagE::erm* | This work |
| BCW72 | *yvhJΩPxylA-mazF (cat)* | This work |
| BCW77 | *ywtFΩPxylA-mazF (cat)* | This work |
| BCW78 | *ywtFΩmsfGFP-ywtF* | This work |
| BCW79 | *yvhJΩmsfGFP-yvhJ* | This work |
| BCW80 | *lytRΩPxylA-mazF (cat)* | This work |
| BCW81 | *lytRΩmsfGFP-lytR* | This work |
| BCW82 | *tagO::erm, ycgO::PxylA-tagO, amyE::Pspac-gfp-mreB (spec), dacA::kan* | This work |
| BDR2061 | *amyE::PxylA-gfp-mbl (spec), mblΩpMUTIN4 (erm)* | (Carballido-Lopez and Errington, 2003) |
| BEB1451 | *hisA1 argC4 metC3 tagO::erm* | (D'Elia et al., 2006) |
| BJS18 | *amyE::PxylA-gfp-mbl (spec)* | (Defeu Soufo and Graumann, 2004) |
| BMD61 | *mblΩmbl-msfGFP (spec)* | This work |
| BRB785 | *yhdG::Pspank-pbpA (phleo), pbpH::spec, pbpA::erm, mblΩPxylA-gfp-mbl (cat)* | (Garner et al., 2011) |
| BEG202 | *ΔtagO::erm amyE::Pxyl-gfp-mbl (spec)* | (Kawai et al., 2011) |
| BEG281 | *ycgO::PxylA-tagO* | This work |
| BEG291 | *tagO::erm, ycgO::PxylA-tagO,* | This work |
| BEG275 | *amyE::Pspac-gfp-mreB (spec)* | (Billings et al., 2014) |
| BEG300 | *tagO::erm, ycgO::PxylA-tagO, amyE::Pspac-gfp-mreB (spec),* | This work |
| BRB4282 | *168 trpC2 ΔtagO::erm* | (D'Elia et al., 2006) |
| bAB343 | *ftsZΩmNeonGreen-15aa-ftsZ, amyE::spc-Pspank-mciZ, ycgO::cat-Pxyl-tagO, tagO::erm* | This work |
| bAB327 | *ftsZΩmNeonGreen-15aa-ftsZ, amyE::Phyperspank-minCD, ycgO::Pxyl-tagO, tagO::erm* | This work |
| bAB388 | *ftsZΩmNeonGreen-15aa-ftsZ, amyE::Physpank-ftsA ycgO::cat-Pxyl-tagO, tagO::erm* | This work |

**Table S3:**
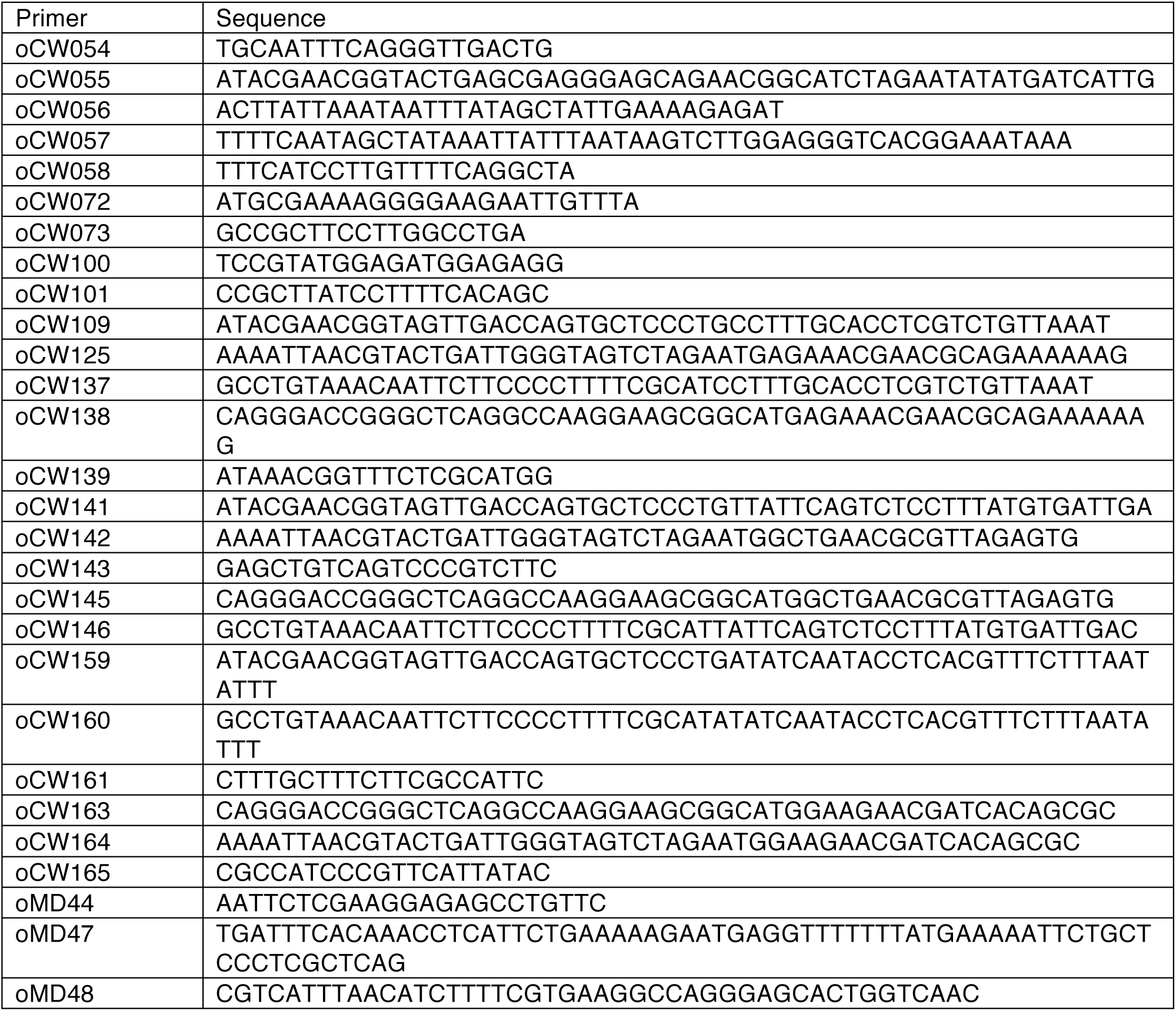
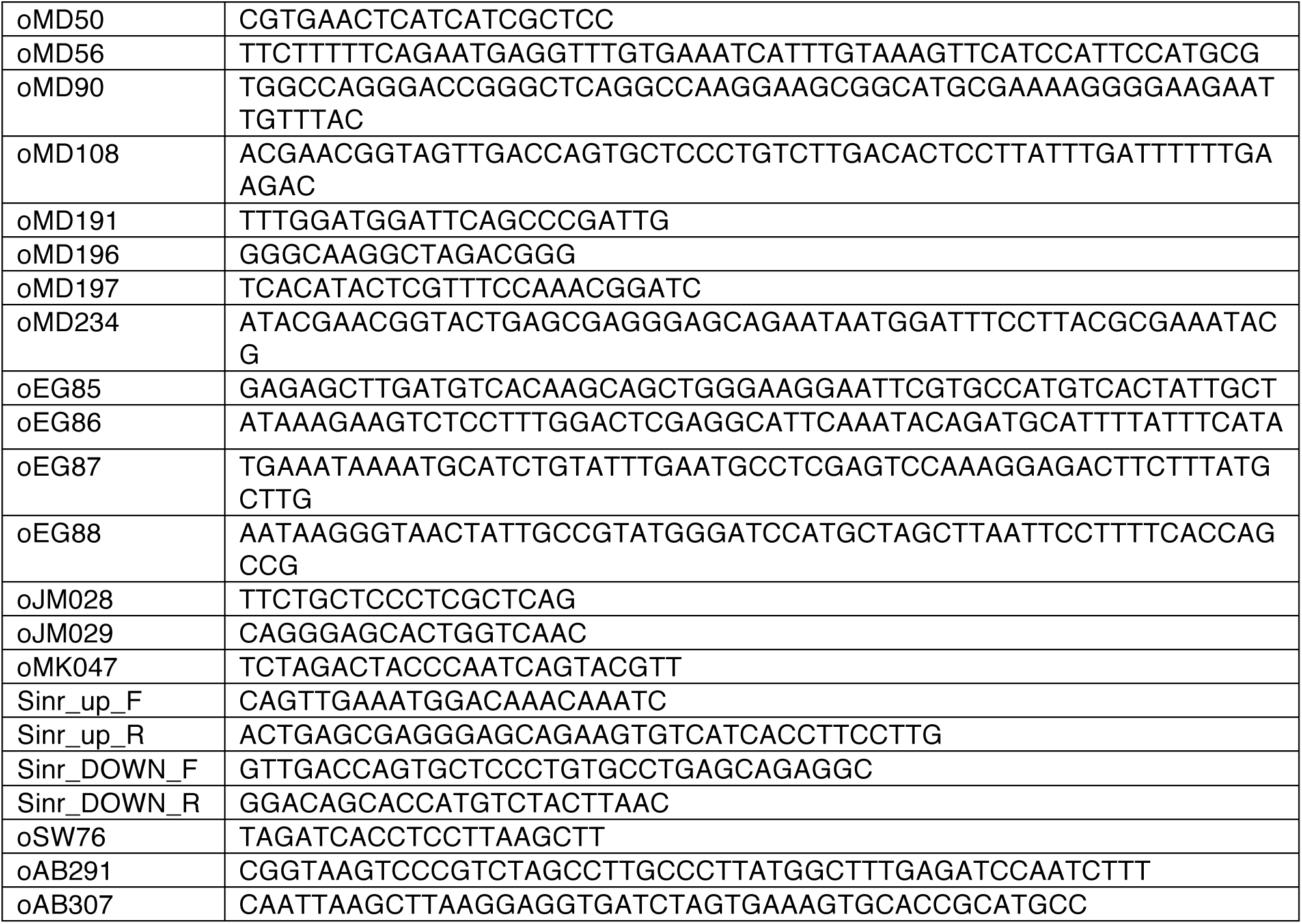
Strains used in this study.

## References

Amir, A., and Nelson, D.R. (2012). Dislocation-mediated growth of bacterial cell walls. Proc Natl Acad Sci USA 109, 9833–9838.

Atilano, M.L., Pereira, P.M., Yates, J., Reed, P., Veiga, H., Pinho, M.G., and Filipe, S.R. (2010). Teichoic acids are temporal and spatial regulators. Proc Natl Acad Sci USA 107, 18991–18996.

Balyuzi, H., Reaveley, D.A., and Burge, R.E. (1972). X-ray diffraction studies of cell walls and peptidoglycans from Gram-positive bacteria. Nature.

Baskin, T.I. (2005). Anisotropic expansion of the plant cell wall. Annu. Rev. Cell Dev. Biol. 21, 203–222.

Billings, G., Ouzounov, N., Ursell, T., Desmarais, S.M., Shaevitz, J., Gitai, Z., and Huang, K.C. (2014). De novo morphogenesis in L-forms via geometric control of cell growth. Mol Microbiol 93, 883–896.

Bonazzi, D., Julien, J.-D., Romao, M., Seddiki, R., Piel, M., Boudaoud, A., and Minc, N. (2014). Symmetry breaking in spore germination relies on an interplay between polar cap stability and spore wall mechanics. Dev. Cell 28, 534–546.

Bringmann, M., Landrein, B., Schudoma, C., Hamant, O., Hauser, M.-T., and Persson, (2012). Cracking the elusive alignment hypothesis: the microtubule-cellulose synthase nexus unraveled. Trends Plant Sci. 17, 666–674.

Chang, F., and Huang, K.C. (2014). How and why cells grow as rods. BMC Biol. 12, 54.

Cho, H., Wivagg, C.N., Kapoor, M., Barry, Z., Rohs, P.D.A., Suh, H., Marto, J.A., Garner, E.C., and Bernhardt, T.G. (2016). Bacterial cell wall biogenesis is mediated by SEDS and PBP polymerase families functioning semi-autonomously. Nat Microbiol 1, 16172.

D'Elia, M.A., Millar, K.E., Beveridge, T.J., and Brown, E.D. (2006). Wall Teichoic Acid Polymers Are Dispensable for Cell Viability in Bacillus subtilis. Journal of Bacteriology 188, 8313–8316.

Defeu Soufo, H.J., and Graumann, P.L. (2006). Dynamic localization and interaction with other Bacillus subtilis actin-like proteins are important for the function of MreB. Mol Microbiol 62, 1340–1356.

Dempwolff, F., Reimold, C., Reth, M., and Graumann, P.L. (2011). Bacillus subtilis MreB orthologs self-organize into filamentous structures underneath the cell membrane in a heterologous cell system. PLOS ONE 6, e27035.

Domĺnguez-Escobar, J., Chastanet, A., Crevenna, A.H., Fromion, V., Wedlich-Söldner, R., and Carballido-Lopez, R. (2011). Processive movement of MreB-associated cell wall biosynthetic complexes in bacteria. Science 333, 225–228.

Gan, L., Chen, S., and Jensen, G.J. (2008). Molecular organization of Gram-negative peptidoglycan. Proc Natl Acad Sci USA 105, 18953–18957.

Garner, E.C., Bernard, R., Wang, W., Zhuang, X., Rudner, D.Z., and Mitchison, T. (2011). Coupled, circumferential motions of the cell wall synthesis machinery and MreB filaments in B. subtilis. Science 333, 222–225.

Harris, L.K., Dye, N.A., and Theriot, J.A. (2014). A Caulobacter MreB mutant with irregular cell shape exhibits compensatory widening to maintain a preferred surface area to volume ratio. Mol Microbiol 94, 988–1005.

Hayhurst, E.J., Kailas, L., Hobbs, J.K., and Foster, S.J. (2008). Cell wall peptidoglycan architecture in Bacillus subtilis. Proc Natl Acad Sci USA 105, 14603–14608.

Holtje, J. (1998). Growth of the stress-bearing and shape-maintaining murein sacculus of Escherichia coli. Microbiol Mol Biol Rev 62, 181–203.

Ishii, T., Matsunaga, T., and Hayashi, N. (2001). Formation of Rhamnogalacturonan II-Borate Dimer in Pectin Determines Cell Wall Thickness of Pumpkin Tissue. Plant Physiol. 126, 1698–1705.

Jain, I.H., Vijayan, V., and O'Shea, E.K. (2012). Spatial ordering of chromosomes enhances the fidelity of chromosome partitioning in cyanobacteria. Proc Natl Acad Sci USA 109, 13638–13643.

Jones, L.J., Carballido-López, R., and Errington, J. (2001). Control of cell shape in bacteria: helical, actin-like filaments in Bacillus subtilis. Cell 104, 913–922.

Kasahara, J., Kiriyama, Y., Miyashita, M., Kondo, T., Yamada, T., Yazawa, K., Yoshikawa, R., and Yamamoto, H. (2016). Teichoic Acid Polymers Affect Expression and Localization of dl-Endopeptidase LytE Required for Lateral Cell Wall Hydrolysis in Bacillus subtilis. Journal of Bacteriology 198, 1585–1594.

Kawai, Y., Marles-Wright, J., Cleverley, R.M., Emmins, R., Ishikawa, S., Kuwano, M., Heinz, N., Bui, N.K., Hoyland, C.N., Ogasawara, N., et al. (2011). A widespread family of bacterial cell wall assembly proteins. Embo J. 30, 4931–4941.

Kawai, Y., Mercier, R., and Errington, J. (2014). Bacterial cell morphogenesis does not require a preexisting template structure. Curr. Biol. 24, 863–867.

Loskill, P., Pereira, P.M., Jung, P., Bischoff, M., Herrmann, M., Pinho, M.G., and Jacobs, K. (2014). Reduction of the peptidoglycan crosslinking causes a decrease in stiffness of the Staphylococcus aureus cell envelope. Biophys J 107, 1082–1089.

Matias, V.R.F., and Beveridge, T.J. (2005). Cryo-electron microscopy reveals native polymeric cell wall structure in Bacillus subtilis 168 and the existence of a periplasmic space. Mol Microbiol 56, 240–251.

Mercier, R., Kawai, Y., and Errington, J. (2013). Excess membrane synthesis drives a primitive mode of cell proliferation. Cell 152, 997–1007.

Olshausen, P.V., Defeu Soufo, H.J., Wicker, K., Heintzmann, R., Graumann, P.L., and Rohrbach, A. (2013). Superresolution imaging of dynamic MreB filaments in B. subtilis--a multiple-motor-driven transport? Biophys J 105, 1171–1181.

Ouzounov, N., Nguyen, J.P., Bratton, B.P., Jacobowitz, D., Gitai, Z., and Shaevitz, J.W. (2016). MreB Orientation Correlates with Cell Diameter in Escherichia coli. Biophys J 111, 1035–1043.

Pandey, R., Beek, Ter, A., Vischer, N.O.E., Smelt, J.P.P.M., Brul, S., and Manders, E.M.M. (2013). Live cell imaging of germination and outgrowth of individual bacillus subtilis spores; the effect of heat stress quantitatively analyzed with SporeTracker. PLOS ONE 8, e58972.

Paredez, A.R., Somerville, C.R., and Ehrhardt, D.W. (2006). Visualization of cellulose synthase demonstrates functional association with microtubules. Science 312, 1491– 1495.

Pooley, H.M., Abellan, F.-X., and Karamata, D. (1993). Wall Teichoic Acid, Peptidoglycan Synthesis and Morphogenesis in Bacillus Subtilis. In Bacterial Growth and Lysis, (Boston, MA: Springer, Boston, MA), pp. 385–392.

Redgwell, R.J., MacRae, E., Hallett, I., Fischer, M., and Perry, J. (1997). In vivo and in vitro swelling of cell walls during fruit ripening. Planta.

Renner, L.D., Eswaramoorthy, P., Ramamurthi, K.S., and Weibel, D.B. (2013). Studying biomolecule localization by engineering bacterial cell wall curvature. PLOS ONE 8, e84143.

Salje, J., van den Ent, F., de Boer, P., and Löwe, J. (2011). Direct membrane binding by bacterial actin MreB. Mol Cell 43, 478–487.

Soufo, H.J.D., and Graumann, P.L. (2010). Bacillus subtilis MreB paralogues have different filament architectures and lead to shape remodelling of a heterologous cell system. Mol Microbiol 78, 1145–1158.

Thomas, K.J., and Rice, C.V. (2014). Revised model of calcium and magnesium binding to the bacterial cell wall. Biometals 27, 1361–1370.

Ursell, T.S., Nguyen, J., Monds, R.D., Colavin, A., Billings, G., Ouzounov, N., Gitai, Z., Shaevitz, J.W., and Huang, K.C. (2014). Rod-like bacterial shape is maintained by feedback between cell curvature and cytoskeletal localization. Proc Natl Acad Sci USA 111, E1025–E1034.

van den Ent, F., Amos, L., and Löwe, J. (2001). Bacterial ancestry of actin and tubulin. Curr. Opin. Microbiol. 4, 634–638.

van den Ent, F., Izoré, T., Bharat, T.A., Johnson, C.M., and Löwe, J. (2014). Bacterial actin MreB forms antiparallel double filaments. Elife 3, e02634.

van Teeffelen, S., Wang, S., Furchtgott, L., Huang, K.C., Wingreen, N.S., Shaevitz, J.W., and Gitai, Z. (2011). The bacterial actin MreB rotates, and rotation depends on cell-wall assembly. Proc Natl Acad Sci USA 108, 15822–15827.

Verwer, R.W., Beachey, E.H., Keck, W., Stoub, A.M., and Poldermans, J.E. (1980). Oriented fragmentation of Escherichia coli sacculi by sonication. Journal of Bacteriology 141, 327–332.

Wachi, M., and Matsuhashi, M. (1989). Negative control of cell division by mreB, a gene that functions in determining the rod shape of Escherichia coli cells. Journal of Bacteriology 171, 3123–3127.

Wang, S., and Wingreen, N.S. (2013). Cell shape can mediate the spatial organization of the bacterial cytoskeleton. Biophysical Journal 104, 541–552.

Wyrick, P.B., and Rogers, H.J. (1973). Isolation and characterization of cell wall-defective variants of Bacillus subtilis and Bacillus licheniformis. Journal of Bacteriology 116, 456–465.

Yao, X., Jericho, M., Pink, D., and Beveridge, T. (1999). Thickness and elasticity of gram-negative murein sacculi measured by atomic force microscopy. Journal of Bacteriology 181, 6865–6875.

## Supplemental References

Agulleiro, J.I., and Fernandez, J.J. (2011). Fast tomographic reconstruction on multicore computers. Bioinformatics 27, 582–583.

Allison, S.E., D'Elia, M.A., Arar, S., Monteiro, M.A., and Brown, E.D. (2011). Studies of the genetics, function, and kinetic mechanism of TagE, the wall teichoic acid glycosyltransferase in Bacillus subtilis 168. J Biol Chem 286, 23708–23716.

Amat, F., Moussavi, F., Comolli, L.R., Elidan, G., Downing, K.H., and Horowitz, M. (2008). Markov random field based automatic image alignment for electron tomography. J. Struct. Biol. 161, 260–275.

Atrih, A., Bacher, G., Allmaier, G., Williamson, M.P., and Foster, S.J. (1999). Analysis of peptidoglycan structure from vegetative cells of Bacillus subtilis 168 and role of PBP 5 in peptidoglycan maturation. Journal of Bacteriology 181, 3956–3966.

Birdsell, D.C., Doyle, R.J., and Morgenstern, M. (1975). Organization of teichoic acid in the cell wall of Bacillus subtilis. Journal of Bacteriology 121, 726–734.

Bisson-Filho, A.W., Hsu, Y.-P., Squyres, G.R., Kuru, E., Wu, F., Jukes, C., Sun, Y., Dekker, C., Holden, S., VanNieuwenhze, M.S., et al. (2017). Treadmilling by FtsZ filaments drives peptidoglycan synthesis and bacterial cell division. Science 355, 739–743.

Carballido-Lopez, R., and Errington, J. (2003). The bacterial cytoskeleton: in vivo dynamics of the actin-like protein Mbl of Bacillus subtilis. Dev. Cell 4, 19–28.

Defeu Soufo, H.J., and Graumann, P.L. (2004). Dynamic movement of actin-like proteins within bacterial cells. EMBO Rep. 5, 789–794.

DeLano, W.L. (2002). The PyMOL molecular graphics system.

Deng, Y., Sun, M., and Shaevitz, J.W. (2011). Direct measurement of cell wall stress stiffening and turgor pressure in live bacterial cells. Phys. Rev. Lett. 107, 158101.

Doyle, R.J., McDannel, M.L., Helman, J.R., and Streips, U.N. (1975). Distribution of teichoic acid in the cell wall of Bacillus subtilis. Journal of Bacteriology 122, 152–158.

Glauner, B., Holtje, J.V., and Schwarz, H. (1988). The composition of the murein of Escherichia coli. J Biol Chem 263, 10088–10095.

Jaqaman, K., Loerke, D., Mettlen, M., Kuwata, H., Grinstein, S., Schmid, S.L., and Danuser, G. (2008). Robust single-particle tracking in live-cell time-lapse sequences. Nat. Methods 5, 695–702.

Kojima, H., Ishijima, A., and Yanagida, T. (1994). Direct measurement of stiffness of single actin filaments with and without tropomyosin by in vitro nanomanipulation. Proc. Natl. Acad. Sci. U.S.a. 91, 12962–12966.

Kremer, J.R., Mastronarde, D.N., and McIntosh, J.R. (1996). Computer visualization of three-dimensional image data using IMOD. J. Struct. Biol. 116, 71–76.

Kuru, E., Hughes, H.V., Brown, P.J., Hall, E., Tekkam, S., Cava, F., de Pedro, M.A., Brun, Y.V., and VanNieuwenhze, M.S. (2012). In Situ probing of newly synthesized peptidoglycan in live bacteria with fluorescent D-amino acids. Angew. Chem. Int. Ed. Engl. 51, 12519–12523.

Kühner, D., Stahl, M., Demircioglu, D.D., and Bertsche, U. (2014). From cells to muropeptide structures in 24 h: peptidoglycan mapping by UPLC-MS. Sci Rep 4, 7494.

Landau, L.D., and Lifshitz, E. (1970). Theory of Elasticity (Pergamon Press).

Mastronarde, D.N. (2005). Automated electron microscope tomography using robust prediction of specimen movements. J. Struct. Biol. 152, 36–51.

Norman, T.M., Lord, N.D., Paulsson, J., and Losick, R. (2013). Memory and modularity in cell-fate decision making. Nature 503, 481–486.

Pettersen, E.F., Goddard, T.D., Huang, C.C., Couch, G.S., Greenblatt, D.M., Meng, E.C., and Ferrin, T.E. (2004). UCSF Chimera—A visualization system for exploratory research and analysis. Journal of Computational Chemistry 25, 1605–1612.

Phillips, R., Kondev, J., Theriot, J., and Garcia, H. (2012). Physical Biology of the Cell (Garland Science).

Safran, S.A. (2003). Statistical thermodynamics of surfaces, interfaces, and membranes (Westview Press).

Schindelin, J., Arganda-Carreras, I., Frise, E., Kaynig, V., Longair, M., Pietzsch, T., Preibisch, S., Rueden, C., Saalfeld, S., Schmid, B., et al. (2012). Fiji: an open-source platform for biological-image analysis. Nat. Methods 9, 676–682.

Schneider, C.A., Rasband, W.S., and Eliceiri, K.W. (2012). NIH Image to ImageJ: 25 years of image analysis. Nat. Methods 9, 671–675.

Sochacki, K.A., Shkel, I.A., Record, M.T., and Weisshaar, J.C. (2011). Protein diffusion in the periplasm of E. coli under osmotic stress. Biophys J 100, 22–31.

Szwedziak, P., Wang, Q., Bharat, T.A.M., Tsim, M., and Löwe, J. (2014). Architecture of the ring formed by the tubulin homologue FtsZ in bacterial cell division. Elife 3, e04601.

Tinevez, J.-Y., Perry, N., Schindelin, J., Hoopes, G.M., Reynolds, G.D., Laplantine, E.,Bednarek, S.Y., Shorte, S.L., and Eliceiri, K.W. (2017). TrackMate: An open and extensible platform for single-particle tracking. Methods 115, 80–90.

Ursell, T., Lee, T.K., Shiomi, D., Shi, H., Tropini, C., Monds, R.D., Colavin, A., Billings, G., Bhaya-Grossman, I., Broxton, M., et al. (2017). Rapid, precise quantification of bacterial cellula dimensions across a genomic-scale knockout library. BMC Biol. 15, 17.

Van den Ent, F., Izoré, T., Bharat, T.A., Johnson, C.M., and Löwe, J. (2014). Bacterial actin MreB forms antiparallel double filaments. Elife 3, e02634.

Whatmore, A.M., and Reed, R.H. (1990). Determination of turgor pressure in Bacillus subtilis: a possible role for K+ in turgor regulation. J. Gen. Microbiol. 136, 2521–2526.

Yu, H., Yan, X., Shen, W., Shen, Y., and Zhang, J. (2010). Efficient and precise construction of markerless manipulations in the Bacillus subtilis genome. Journal of Microbiology

Zhong-Can, O.Y., and Helfrich, W. (1989). Bending energy of vesicle membranes: General expressions for the first, second, and third variation of the shape energy and applications to spheres and cylinders. Phys. Rev. A 39, 5280–5288.

